# Active sensing explains cortical variability and improves visual information

**DOI:** 10.64898/2026.10.12.758571

**Authors:** Ryan A. Ressmeyer, Declan P. Rowley, Tejas Kothapalli, Dekel Galor, Ruei-Jr Wu, Michele Rucci, Jude F. Mitchell, Daniel Butts, Alexander C. Huk, Jacob L. Yates

## Abstract

Active vision is often considered at the scale of saccades, with each fixation treated as a static visual sample. Yet the eyes continue to move during fixation, continually transforming the retinal input. Here we measured these movements at high precision while recording neuronal populations in the foveal primary visual cortex of awake marmosets viewing repeated flashed natural images. Fixational eye movements accounted for most conditional firing-rate variability and nearly all shared variability not explained by the trial-averaged response. An image-computable model captured this modulation largely from gaze-dependent retinal input; explicit eye-position and eye-velocity signals added little predictive power. Under natural viewing, retinal motion redistributed image power across temporal frequencies, and motion effects grew when this power engaged spatiotemporal tuning. In model-derived populations, fixational motion increased the spatial information carried per spike relative to stabilized input, with sharpening growing across successive nonlinear stages. Thus, activity commonly assigned to cortical noise is not just structured visual reafference generated by the movement of the observer, but is beneficial signal which sharpens spatial representations.

## Introduction

In natural settings, sensation is inseparable from action. An animal’s own movements continually shape the signals arriving at its sensory receptors. Neural activity therefore depends not only on external stimuli but also on the animal’s recent movement history. Such sensory consequences of movements are termed “reafference”. Vision provides an unusually precise example of this coupling between sensation and action. The eyes are never at rest [1, 2], and fixational eye movements (FEMs)—including drift and microsaccades—continually translate the image across the retina. Decades of psychophysical work show that these minute displacements support fine spatial vision [3–5]. Retinal motion converts spatial structure into temporal modulation. Theoretical work treats this temporal signal as a carrier of spatial information [6–8] that can be read out to resolve detail finer than the cone mosaic alone would permit [9, 10]. Consistent with this account, neurophysiological studies have found substantial effects of FEMs in the retina [11] and primary visual cortex (V1) [12–15]. Yet how this self-generated retinal motion shapes visual encoding across cortical populations remains unknown.

Answering this question requires determining how much cortical activity varies with FEMs and how those variations relate to the response shared across repeated presentations of the same displayed stimulus. Conventional analyses often assume that these presentations deliver the same sensory input, attributing trial-to-trial differences in response to internally generated variability [16, 17]. During fixation, this assumption can fail, because FEMs dynamically change the retinal image even when the displayed stimulus is repeated [15]. Analyses that do not account for changes in the eye’s position therefore risk attributing differences in retinal input to intrinsic variability of the neural circuit. Separating these components is necessary to interpret estimates of response reliability and the mechanisms underlying that reliability [16–19].

The effects of FEMs are expected to be especially consequential for foveal vision. The primate fovea subtends the central ~5–8 degrees of visual angle across species [20], supports the highest spatial acuity, and is disproportionately represented in V1 and other cortical areas [21], with roughly one-third of marmoset V1 devoted to foveal vision [22]. FEMs can displace the retinal image by amounts comparable to the receptive-field sizes of individual foveal neurons. To understand how these displacements shape the cortical activity supporting fine spatial vision, we used laminar electrode probes to record spiking activity from ensembles of neurons in the foveal representation of marmoset V1 during alert fixation of naturalistic images. Relating this activity to retinal input required measuring eye position with high precision, which we achieved using a custom digital dual Purkinje image eye tracker [23–25]. We repeatedly presented a fixed sequence of naturalistic images while recording cortical responses and the eye’s trajectory on each presentation. Comparing neural responses with respect to the similarity of the eye’s trajectory between repeats allowed us to separate trial-to-trial variability associated with FEMs from variability that remained unexplained.

FEMs accounted for approximately 70% of the conditional firing-rate variance in individual neurons, exceeding the contribution captured by the trial-averaged response. Accounting for the effects of FEMs also substantially reduced conventional estimates of residual variability (“noise”), both independent and shared between neurons. FEM-driven variance was concentrated in a few dominant population dimensions that overlapped those of the trial-averaged stimulus-driven response. A deep neural network encoding model predicted much of the single-trial modulation from retinal input, leaving a much smaller additional contribution from explicit eye-position and eye-velocity inputs, suggesting that feedforward retinal processing is sufficient to account for the dependence of V1 activity on FEMs.

In further model-based analyses, FEMs increased spatial information in select neural subpopulations compared with stabilized retinal input. These effects depended on eye-movement dynamics, stimulus content, and neuronal spatiotemporal tuning. Combining contributions from successive nonlinear processing stages increased spatial sharpening in the model, suggesting a nonlinear contribution to the information improvements. Together, these results demonstrate the fundamental role of active sensing in foveal vision, whereby fine-scale eye movements drive large changes in cortical ensemble activity while sharpening the representation of spatial features. Reafference has been demonstrated as a key element of active sensing in other sensory modalities; our results show that it also plays a central role in the highest-acuity form of vision.

## Results

### Fixational eye movements exert large effects on the structure of neural responses in foveal V1

We recorded from populations of 10–118 analysis-eligible units per session (1,022 units across 19 sessions) in the foveal representation of the primary visual cortex (V1) in two awake marmosets. Subjects were trained to maintain stable fixation within a 2° window while viewing a repeated sequence of rapidly flashed, whitened natural images presented at 20 Hz (Fig. 1A). Eye position was monitored with a high-precision digital dual Purkinje image eye tracker [23–25] and typically remained well within the fixation window across trials (Fig. 1B). The distributions of eye position during fixation were similar to those observed in humans and macaques [25, 26]. Receptive fields of recorded units were confined to eccentricities of *<*1° (Fig. 1C). As expected, activity was elicited in the recorded units by the visual stimuli presented within their receptive fields (RF), and the resulting patterns of spikes displayed variability between trials (Fig. 1D).

**Figure 1.**
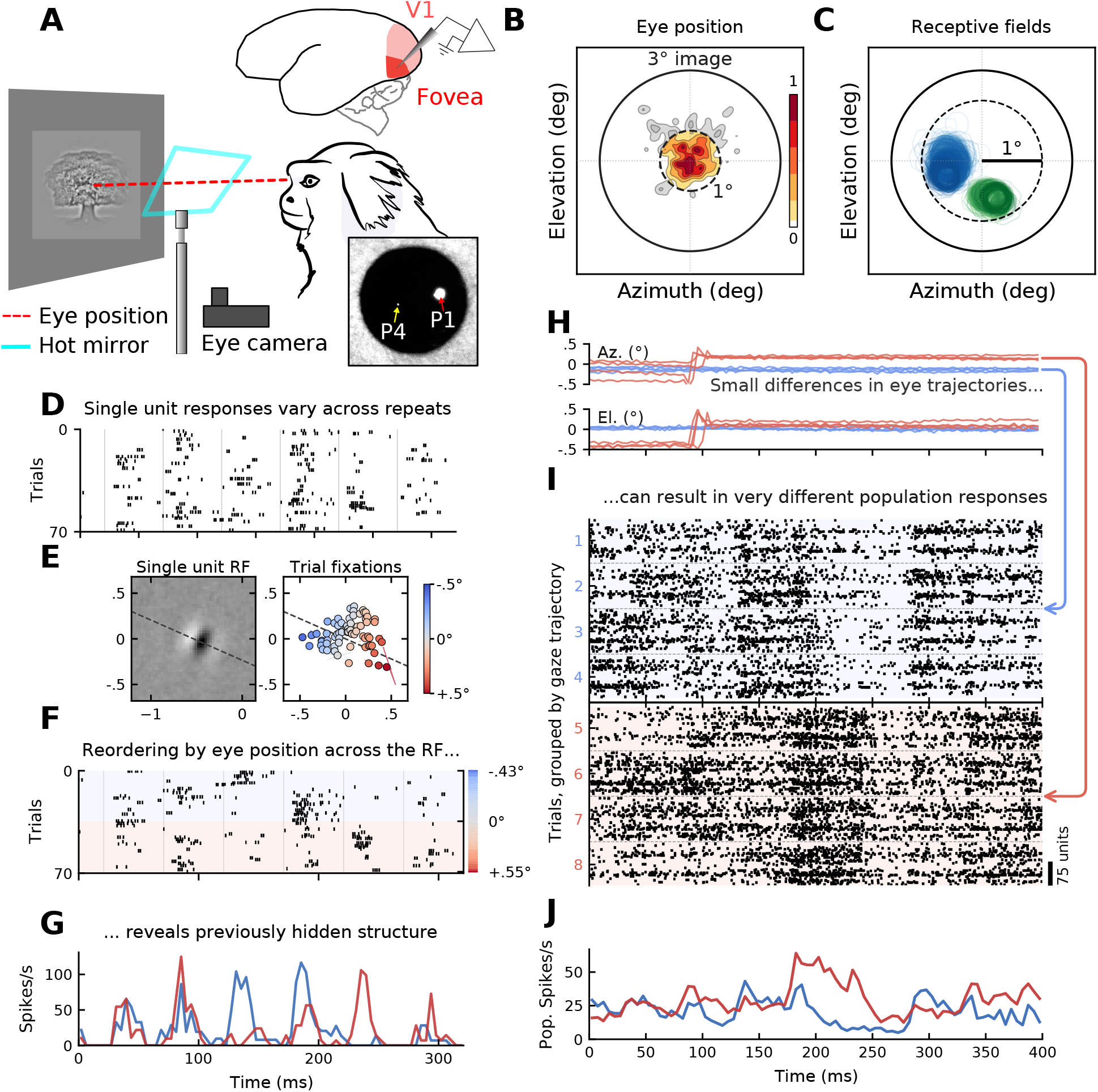
Small eye movements structure neural activity within the foveal representation of the primary visual cortex. **(A)** Schematic of the experimental paradigm. A head-fixed marmoset fixated a rapidly updating sequence of flashed images (3° diameter). Electrical activity was recorded using laminar probes from the lateral surface of V1. The eye’s position was tracked using a high-precision digital dual Purkinje image eye tracker [23–25]. The eye was imaged through a hot mirror that allowed the subject to view the stimulus. A representative image of the Purkinje images is shown in the inset. **(B)** Distribution of eye positions during fixation for a representative experiment. Marmosets maintained tight fixational control, which was further refined by only considering eye positions within a 1° diameter circle for analysis. **(C)** Receptive field (RF) locations. The extent of each unit’s RF is drawn as a contour. Blue contours represent an ensemble of units recorded from monkey A; green contours show an example ensemble from monkey L. All units had receptive fields within 1° from the fovea. **(D)** Trial rasters for the example unit, ordered by trial number. The single-unit responses vary across repeated stimulus presentations. **(E)** Left: The RF for the unit shown in (D), which was obtained by spike-triggered averaging on a separate flashed gabor noise stimulus [23]. The RF displays clear, Gabor-like structure. Right: The average eye position in a 50 ms window across trials. Points are colored according to their projection onto the line orthogonal to the subunits of the RF. **(F)** Trial rasters shown in (D), now ordered by the projection of the mean eye position across the RF in 50 ms windows. Vertical gray lines denote the window boundaries. Smooth variation in trial-to-trial responses emerges when sorting by eye position, indicating a relation between the eye’s position and the neural response. **(G)** The peristimulus time histogram of responses with positive projections across the RF (blue) and negative projections (red). Average stimulus-evoked responses diverge strongly between 100 ms and 250 ms. **(H)** Eye position over time for eight representative trials colored in red and blue. The eye position traces are highly self-similar within groups, but not between groups. **(I)** Population rasters for the eight representative trials from (H). There is clear similarity between population responses on trials with similar eye trajectories, and large differences between trials with dissimilar eye trajectory, despite the fact that the difference in position is just fractions of a degree. **(J)** Spiking activity averaged across all recorded units for the four red trials and four blue trials over the same time course. The average population firing rate diverges strongly around 200 ms in the trial.

Trial-to-trial variability was strongly related to the fine-scale position of the eye. For each unit we measured the displacement of the eye between trials, projected onto the axis across which the unit’s receptive field is tuned (Fig. 1E, Extended Data Figure 1). Sorting the stimulus-evoked responses by this displacement revealed clear structure in the trial-to-trial fluctuations (Fig. 1F), and splitting trials by the sign of the projection produced markedly different response timecourses (Fig. 1G). FEMs can therefore substantially structure the activity of individual units.

Response structuring by FEMs was evident at the population level as well. We visualized population level rasters from multiple trials while conditioning on clusters of similar gaze trajectories. We identified two groups of trials with highly self-similar eye trajectories, but which differed from one other (Fig. 1H). Population responses were reproducible within each group (trials 1-4, blue), whereas trials with distinct trajectories exhibited systematically different patterns of activity (trials 5-8, red; Fig. 1I, Extended Data Figure 2). These distinct profiles are apparent in the group-averaged population responses (Fig. 1J)– even though the stimulus was identical across all trials.

These visualizations make a qualitative case that FEMs structure responses at both the single-unit and population levels. A comprehensive and quantitative account requires additional statistical dissection. A simple, intuitive approach would select a subset of trials on which the eye followed nearly identical trajectories. However, such trials are rare. We therefore turned to a statistical approach that quantifies how FEMs modulate population variability without requiring perfectly-matched repetitions [15].

### Active sensing accounts for most residual neural variability

To quantify the contribution of FEMs to neural variability, we extended a recently developed method for variability decomposition ([15]; Methods). Because the same time-varying image was presented on every trial, any two trials share the same stimulus at every given timepoint, and differ only in the eye’s position during the preceding moments. The similarity of their recent eye trajectories therefore determines whether the two spike counts arose from the same retinal input (Fig. 2A). We therefore estimated variability from only those trial pairs whose recent trajectories fell within a given distance of one another, and varied that distance (Fig. 2B). When the distance is large enough to admit every pair, the estimate recovers the eye-position-blind residual, which is the variability that remains once the trial-averaged response is subtracted. As the distance shrinks, only pairs whose eye positions nearly coincided over the preceding moments contribute, and the estimate falls toward a floor. The gap between the uncorrected residual and that floor is the variability driven by fixational eye movements, and the floor is the residual variability that survives once eye position is accounted for.

**Figure 2.**
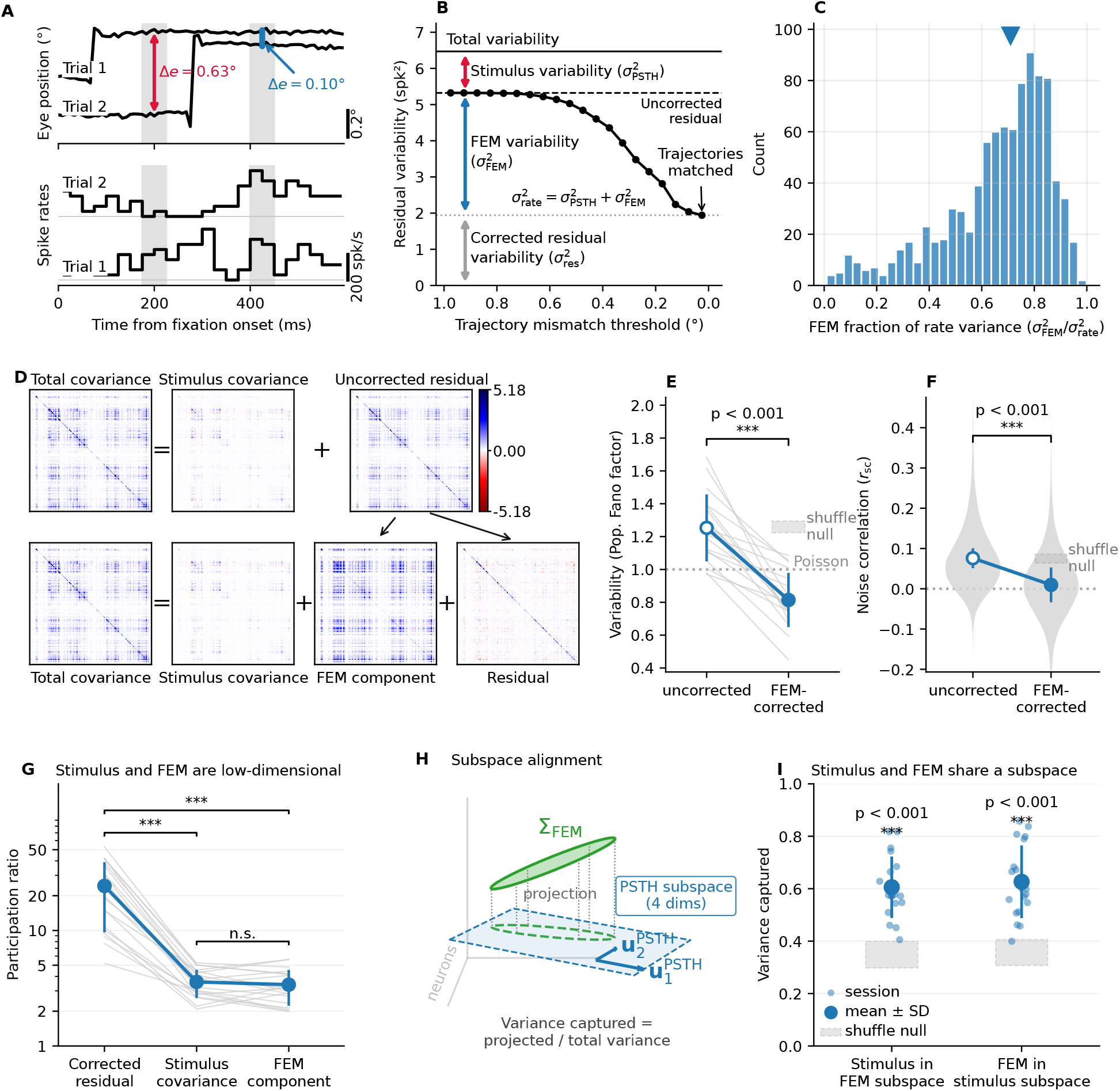
Fixational eye movements dominate single-unit and shared variability, and contribute a compact population covariance component aligned with the stimulus-driven subspace. **(A)** The comparison underlying the decomposition, for an example unit. At a fixed time-point in the frozen stimulus sequence, eye trajectories are compared across trials. Top: eye-position traces for two trials whose eye position is closely matched at one timepoint (blue, Δ*e* = 0.10°) and divergent at another (red, Δ*e* = 0.63°). Bottom: the corresponding per-bin spike rates. Responses are similar when the eye’s position is aligned across trials, while divergent eye positions yield divergent responses despite identical stimulus input. **(B)** Eye-attributable variability for the same unit, recovered by tightening the matching threshold. As trial pairs are required to have increasingly similar trajectories (Δ*e < x*), the estimate falls from the uncorrected residual toward the residual variability that survives once eye position is accounted for. The gap (blue, 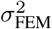) is the conditional rate variance attributable to fixational eye movements. Together with the variance of the trial-averaged response (red, 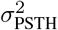), it constitutes the conditional rate variance 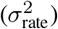. **(C)** Distribution across units of the FEM fraction of single-unit conditional rate variance 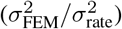; the triangle marks the pooled median. The distribution is biased toward one (median 0.71, 95% CI [0.69, 0.72]), indicating that most trial-to-trial rate variance is attributable to fixational eye movements. **(D)** Covariance decomposition for an example session. Top: the conventional split, total covariance = Σ_PSTH_ + Σ_res,unc_, assigns all covariance not captured by the PSTH to a single component. Bottom: that uncorrected residual is further partitioned into an eye-movement–dependent component Σ_FEM_ and a residual Σ_res_. Off-diagonal structure in the uncorrected residual migrates almost entirely into Σ_FEM_, leaving Σ_res_ near-diagonal. (**E**)Population Fano factor, uncorrected vs. FEM-corrected. Thin grey lines are individual sessions; the marker is the across-session mean ± SD. The Poisson reference (Fano = 1) is dotted and the grey band is the shuffle null. (**F**)Per-pair noise correlation (*r*_sc_), uncorrected vs. FEM-corrected. Grey violins show the pooled per-pair distribution; the marker is the across-session mean ± SD and the grey band the shuffle null. **(G)** Participation ratio of the three covariance components — residual (Σ_res_), stimulus-locked (Σ_PSTH_), and eye-movement–dependent (Σ_FEM_). One faint line per session joins its three values; the marker is the across-session mean ± SD (log scale). Variance in the stimulus and FEM components is concentrated in comparably few dominant dimensions relative to the residual. **(H)** Schematic of the subspace-alignment test. FEM covariance (green disk) is projected orthogonally onto the sub-space spanned by the leading eigenvectors of Σ_PSTH_ (columns of *U*; blue plane). The fraction of variance retained, *f*_FEM→PSTH_ = tr(*U* ^⊤^Σ_FEM_*U*)/tr(Σ_FEM_), measures how much FEM-driven variance lies within these dominant stimulus-locked dimensions. This is one of the two directions shown in (I), and is distinct from the FEM fraction of conditional rate variance in (C). **(I)** Variance captured in each direction — stimulus-locked variance lying in the FEM subspace, and FEM variance lying in the stimulus-locked subspace — with one point per session and the across-session mean ± SD. The grey dashed band is the shuffle null of the mean (the eye-trajectory shuffle). Both directions lie well above the null, indicating substantial overlap between the dominant dimensions of FEM-driven and stimulus-locked population variability. [All panels use a 25 ms counting window. Significance markers (∗) denote comparison against the eye-trajectory shuffle null (E, F, I) or paired tests across components (G); numeric values are reported in the main text.]

We define the conditional rate variance (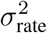 Fig. 2B) as the total variance accounted for by matching trials at the same timepoint on their recent eye trajectories. It combines the variance of the trial-averaged response (i.e., the peri-stimulus time histogram [PSTH]) 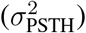and the trial-to-trial variance attributable to differences in the eye’s position 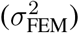. From these, we computed the fraction of the conditional rate variance attributable to fixational eye movements, which had a median of 0.71 across the population (IQR [0.57, 0.81]; *n* = 939 units; Fig. 2C). Neural responses therefore varied more with trial-to-trial differences in eye position than with the stimulus itself, despite fixation behavior that would conventionally be considered as under stringent control.

Conventional measures of neural variability compare each trial to the trial-averaged response, so effects of differences in retinal input are assigned within the conventional, uncorrected residual 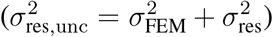. Our decomposition instead estimates the residual after accounting for differences in recent eye position between trials. This partitioning lowered estimates of the population Fano factor from 1.25 to a sub-Poisson value of 0.81 (a reduction absent under a trajectory-shuffle null, empirical *p* ≤ 0.001; Fig. 2E). Units in foveal V1 are therefore substantially less variable than the conventional residual implies.

The decomposition also partitions covariance across simultaneously recorded units into stimulus-locked (Σ_PSTH_), eye-movement–dependent (Σ_FEM_), and residual (Σ_res_) components (Methods; Fig. 2D). Correlations computed from the conventional residual, which combines the latter two components (Σ_res,unc_ = Σ_FEM_ + Σ_res_), are conventionally termed “noise correlations” [27, 28]. The off-diagonal structure of this conventional residual was almost entirely attributable to Σ_FEM_, leaving the FEM-corrected residual (Σ_res_) nearly diagonal. Before FEM correction, noise correlations averaged 0.076 (95% CI [0.065, 0.086]), comparable to previous reports [29, 30]. After correction, they fell to 0.010 (95% CI [−0.010, 0.029]), with a confidence interval that included zero. This reduction exceeded that expected under the trajectory-shuffle null (Δ*r*_sc_ = −0.066, 95% CI [−0.084, 0−.050]; shuffle-null interval [−0.012, 0.010]; empirical *p ≤* 0.001; Fig. 2F). Differences in eye position across trials thus account for almost all of the shared variability left by the trial-averaged response.

FEM-driven variability was also concentrated in a few dominant population dimensions. The participation ratio of Σ_FEM_ averaged 3.4 ± 1.2 across sessions, roughly seven-fold more compact than the residual (24.3 ± 14.7) and not detectably different from the stimulus-locked covariance, Σ_PSTH_ (3.6 ± 1.0; sign test, 12 of 19 sessions, *p* = 0.36; Fig. 2G). With much of the variance in both components concentrated in a few dimensions, those dimensions could overlap or be largely distinct. Both arrangements are plausible. Changes in the displayed image and eye movements both alter the visual features falling within neuronal receptive fields, potentially recruiting similar population response patterns and aligning the dominant dimensions. However, a neuron’s responsiveness across images need not predict its sensitivity to displacements of the stimulus. Small translations may strongly modulate neurons whose receptive-field subregions lie near local image contours while leaving other visually responsive neurons relatively unaffected. FEM-driven variability could therefore occupy dimensions distinct from those dominating the stimulus-locked response. We therefore asked whether the dominant population dimensions of these two covariance components aligned.

We quantified this alignment by computing the fraction of each component’s variance captured by the four leading eigenvectors of the other (Fig. 2H; Methods). Larger fractions indicate greater overlap with the other component’s dominant population dimensions. Because this measure is asymmetric, we evaluated both directions, each against a trajectory-shuffle null that broke the coupling between eye position and population activity. The leading FEM subspace captured a majority of the stimulus-locked variance (0.61 ± 0.12), and the leading stimulus-locked subspace captured a majority of the FEM-driven variance as well (0.63 ± 0.14; means ± SD across sessions). Both fractions exceeded their respective shuffle nulls (≈0.35; *p ≤* 0.001 for both directions; Fig. 2I). FEM-driven variability therefore largely overlaps the dominant subspace spanned by the stimulus-locked population response.

### Retinal input predicts FEM-dependent population activity

The prior analyses show that modulation associated with FEMs overlaps the dominant dimensions of the stimulus-related response, but they do not identify the source of that modulation. Under one account, fixational eye movements displace the image on the retina and influence activity in V1 through primarily feed-forward circuits originating in the photoreceptors (i.e., reafference). Under the other, an internal signal reporting the eye’s position and motion modulates V1 independently of the image on the retina (i.e., extraretinal modulation). Both accounts are consistent with the reported population geometry, but they assign the variability to fundamentally different sources.

We dissociated these two potential accounts statistically by fitting a single image-computable feedforward neural network model to the recorded V1 population. The model was trained to predict responses to gratings, Gabors, and natural images during free viewing [23]. The fixated flashed-image condition analyzed throughout this paper was withheld from training, so the Figure 3 predictive metrics measure generalization across both stimulus and behavioral context (Fig. 3A).

**Figure 3.**
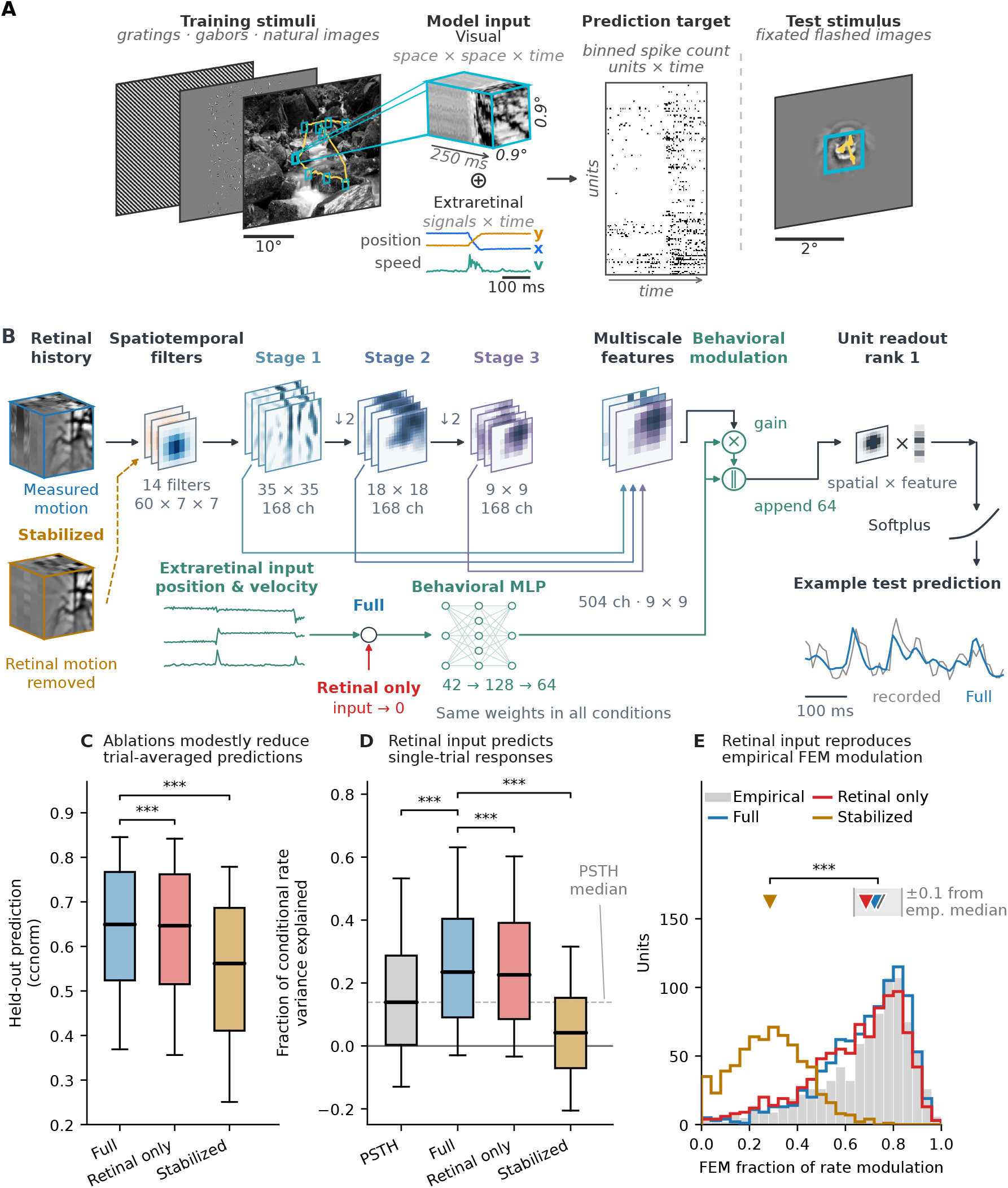
An image-computable model reproduces eye-movement driven variability through feedforward computation on retinal input. **(A)** Stimuli and retinal input. The model is trained on grating, Gabor, and natural image stimuli during free viewing, and evaluated on the held-out flashed image dataset. Its inputs are the gaze-contingent retinal stimulus, a space × space × time reconstruction of the stimulus history accounting for measured eye position, and separate extraretinal eye-position and eye-velocity covariates. **(B)** Model architecture and input interventions. A gaze-contingent retinal movie spanning approximately 0.9°× 0.9° × 250 ms (35 × 35 pixels × 60 lags) passes through 14 learned spatiotemporal filters and three spatial convolutional stages. Features from all three stages are combined into a multiscale representation. A separate pathway transforms eye position and velocity into multiplicative gains on visual features and 64 additional feature channels appended before the readout. Each unit’s readout combines spatial and feature weights, followed by a softplus nonlinearity to predict firing rate. The model shares 3.47 million parameters across 2,790 recorded units from 30 sessions, with 655 readout parameters fitted separately for each unit. The three conditions in C–E use identical fitted weights. Full (blue) retains both inputs; Retinal only (red) zeroes the extraretinal input; Stabilized (ochre) fixes the retinal crop at one session-global gaze position while retaining extraretinal input and stimulus changes. **(C)** Held-out, trial-averaged prediction (noise-normalized correlation, CC_norm_) for the full, retinal-only, and stabilized-retina model conditions (*n* = 962 units across 19 sessions). Relative to the full model, the median paired changes were Δ = −0.005 for retinal-only (0.8% reduction; *p* = 2.4 × 10^−37^) and Δ = −0.082 for stabilized-retina (12.6% reduction; *p* = 9.8 × 10^−69^). Percentages are relative to the full-model median; tests are paired Wilcoxon signed-rank tests. **(D)** Single-trial prediction as a fraction of the conditional rate variance measured in Figure 2. Median paired changes were +0.090 for full minus PSTH (95% CI [+0.068, +0.122]; *p* = 2.0 × 10^−4^), −0.007 for retinal-only minus full (95% CI [−0.011, *−*0.004]; *p* = 2.0 × 10^−4^), and −0.174 for stabilized-retina minus full (95% CI [−0.203, *−*0.152]; *p* = 2.0 × 10^−4^). Confidence intervals and tests use the session-cluster bootstrap. The full-model gain was 65% of the PSTH median; the stabilization loss was 74% of the full-model median. The dashed line marks the trial-average median and the solid line marks zero captured variance. Values above one were retained because the denominator is an estimate, not a hard upper bound. Brackets in C and D show ^∗^*p <* 0.05, ^∗∗^*p <* 0.01, and ^∗∗∗^*p <* 0.001. **(E)** Distribution of the FEM fraction of conditional rate variance 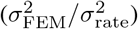 for the recorded units and each model condition. Downward triangles mark distribution medians. The empirical median was 0.736, compared with 0.722 for the full model, 0.686 for the retinal-only model, and 0.285 for the stabilized-retina model. The shaded band shows the empirical median ± 0.1. Paired two-one-sided tests with this equivalence margin supported equivalence for the full and retinal-only models (*p* = 1.3 × 10^−30^ and *p* = 5.9 × 10^−20^, respectively), but not for the stabilized-retina model (*p* = 1.000). The empirical and stabilized-retina estimates differed by a median of 0.423 (paired Wilcoxon *p* = 2.5 × 10^−88^). In C and D, boxes span the interquartile range, median lines mark the medians, and whiskers span the 10th–90th percentiles across units.

The model predicts neural responses from a 250-ms retinal history using a full-history spatiotemporal convolution followed by three spatial convolutional stages and unit-specific readouts (Fig. 3B). The retinal input is gaze-contingent, so fixational eye movements move the image across the model’s input window. A separate extraretinal pathway converts eye position and temporally filtered eye velocity into feature gains and spatially constant additive inputs using a multilayer perceptron model architecture. This is the only route by which eye state can affect the model’s response without changing the retinal input. (“Extraretinal” generically refers to this model pathway; the model does not identify whether an analogous biological signal arises from an efference copy, proprioception, or some other indirect source). Holding the fitted weights fixed, we compared the full model against two ablated versions. In the retinal-only condition, we set the extraretinal input to zero, testing whether that pathway improves prediction once retinal image motion has been specified. In the complementary stabilized-retina condition, we fixed the retinal sampling position for every trial at the center of each session’s gaze distribution, removing the image motion caused by fixational eye movements while the extraretinal input remained intact.

The full and retinal-only models achieved similar trial-averaged accuracy. The full model predicted trial-averaged responses with a median noise-normalized correlation of CC_norm_ = 0.649 across *n* = units from 19 sessions (Fig. 3C). Removing the extraretinal input resulted in only a small performance decrease (median 0.647; median paired ΔCC_norm_ = −0.005; paired signed-rank *p* = 2.4 × 10^−37^). Conversely, stabilizing the retinal input had an order-of-magnitude larger detrimental effect (median 0.562; median paired Δ = −0.082; *p* = 9.8 × 10^−69^). The paired losses were 0.8% and 12.6% of the full-model median, respectively. Much of the trial-averaged response remained predictable, motivating a more stringent assessment at the level of single trials.

We therefore scored each model condition’s prediction on single trials by the fraction of the conditional rate variance measured in Figure 2 that it captured (Methods). This score is zero for a constant-value predictor and one for an ideal predictor of the conditional rate. For comparison, we predicted each trial’s response from the average of the remaining repeats. This leave-one-out trial average estimates the stimulus-locked response that defines the prediction ceiling when trial-to-trial fluctuations are treated as noise [31]. It captured a median of 0.139 of the rate variance, whereas the full model captured 0.235 (median paired Δ = +0.090, 95% CI [+0.068, +0.122]; *n* = units from 19 sessions; session-clustered bootstrap, Methods; Fig. 3D). This amounted to a 65% improvement over the trial-average median, suggesting that the model captured substantial FEM-driven variability around the mean response.

Most of the model’s single-trial predictive advantage persisted when the explicit extraretinal inputs were removed. We found that the extraretinal pathway learned structured modulation linked to saccades (Extended Data Figure 3). However, zeroing the eye position and velocity terms cost very little in terms of captured variance (median 0.226; median paired Δ = −0.007, 95% CI [−0.011, *−*0.004]). In contrast, stabilizing the retinal input reduced the median fraction to 0.042, below the trial-average median (paired Δ = −0.174, 95% CI [−0.203, *−*0.152]; Fig. 3D). This paired loss was 74% of the full-model median. Single-trial prediction therefore depended strongly on gaze-contingent retinal input; removing the separate eye-state input cost substantially less than stabilizing the retinal image.

In an additional analysis, we asked over what timescale reafferent input contributed to accurate model predictions. We compared fixing gaze at one position for the entire session, at each trial’s mean position, or at the latest position within each 250-ms input history. Preserving each history’s final retinal position avoided most of the loss in accuracy caused by fixing gaze across whole trials or sessions. Thus, accurate prediction depended more on tracking moment-to-moment retinal position than on retaining the motion leading to that position (Extended Data Figure 4).

We then asked whether the model reproduced the relative magnitude of FEM-driven modulation measured directly from the recorded units. We applied the same decomposition used in Figure 2 to each condition’s predicted rates and compared the resulting FEM fractions with those estimated from the recorded responses (Methods). The full and retinal-only models produced similar FEM fractions, with medians of 0.722 and 0.686, respectively, compared with an empirical median of 0.736. Paired equivalence tests supported mean empirical-minus-model differences within a ± 0.1 margin (TOST *p* = 1.3 × 10^−30^ and *p* = 5.9 × 10^−20^, respectively; Fig. 3E). In contrast, stabilizing the retinal input reduced the median FEM fraction to 0.285 (median paired empirical-minus-model difference 0.423; paired signed-rank *p* = 2.5 × 10^−88^). Within the fitted model, retinal input alone was sufficient to reproduce the observed FEM fraction, and global stabilization reduced its median by more than half.

These results provide statistical evidence for a substantial reafferent contribution to FEM-dependent activity in foveal V1. Although the model need not reflect the underlying biological circuitry by construction, it does demonstrate that retinal input alone is sufficient to capture the single-trial predictive advantage over the trial average and to reproduce the observed FEM fraction. The small additional predictive contribution of the extraretinal pathway is consistent with extraretinal influences playing a secondary, modulatory role under the conditions of our experiments.

### Fixational eye movements sharpen spatial population codes

The preceding analyses showed that FEMs account for most conditional rate variance in foveal V1 and provided evidence that this modulation is predominantly reafferent. We next asked how the retinal motion driving this variability either benefits or harms the cortical representation of visual features. The recorded data alone could not resolve this question for two reasons. First, any visual feature is represented across the cortical sheet by a population of neurons with varying receptive field locations, while the recordings sample only a subset of neurons in a single cortical column with generally overlapping receptive fields. Second, a direct test of the causal effects of FEMs on visual representations would require matched observations with retinal motion removed, which were unavailable in the recorded datasets. We therefore used the fitted encoding model to make inferences about how FEMs shape V1 representations in complete populations with both observed and counterfactual (i.e., simulated) patterns of eye movements.

In this analysis, we replicated the spatial readout of every fitted unit across the modeled visual field, producing a pseudo-population of identically tuned neurons that differed only in receptive-field position (Fig. 4A). Each activation map gives the predicted firing rate at each receptive-field position, assuming the existence of neighboring unmeasured neurons with identical tuning. We summarized each map by its mean firing rate and its spatial information per predicted spike. The latter, measured using single-spike information (SSI; Methods), quantifies how much spatial information the feature-tuned population conveys per spike, independently of mean firing rate. Their product gives the instantaneous predicted spatial information rate per neuron under this single-spike measure. We compared both quantities between activation maps generated by measured fixation trajectories and those generated by matched movies in which the retinal input was stabilized.

**Figure 4.**
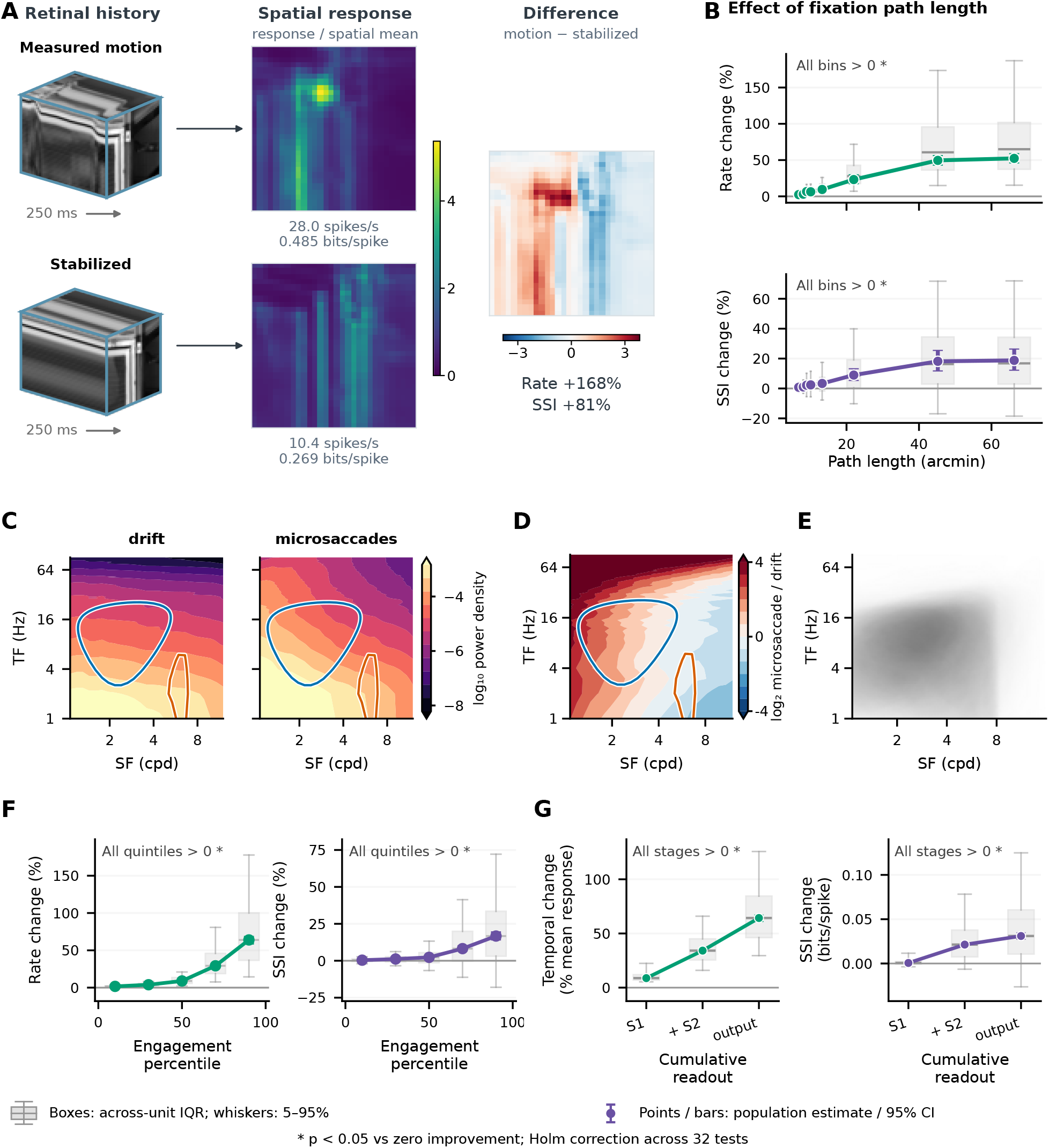
Fixational eye movements engage model-derived passbands and sharpen spatial coding. Fixational eye movements engage model-derived spatiotemporal passbands and sharpen spatial coding in the model. **A**, A filtered, measured eye-movement trace and its stabilized counterpart generate 250-ms natural-image movies matched at the mean first-layer peak lag. Horizontal arrows indicate that each movie passes through the same trained model to generate a spatial activation map for one illustrative unit. Both response maps show response divided by its spatial mean on a shared color scale; the rightmost map shows motion minus stabilized on a symmetric scale. In this example, motion increases firing rate and single-spike information (SSI) and sharpens the normalized spatial response. **B**, Firing-rate and SSI changes versus fixation path length across 725 units. Colored curves show spike-weighted population estimates. **C**, Filled contours of equal-mass natural-image power for 47 drift windows and 18 microsaccade-containing windows. Both conditions use one-second records filtered continuously with a positive Gaussian kernel (*σ* = 6 ms); microsaccade amplitudes are below 1°. Each trace’s nonzero-temporal-frequency power is normalized to unit mass before averaging with equal animal weights. Blue and orange contours show fitted 55%-response passbands for two examples selected from the 145 strictly validated units; the same contours appear in **D. D**, Log_2_ microsaccade/drift power ratio with the example passbands overlaid, showing their overlap with the redistributed retinal power. **E**, Spatiotemporal tuning for all modeled units. Grayscale shading shows population tuning occupancy. **F**, Firing-rate and SSI modulation versus the within-unit percentile of motion-induced tuning-relevant power. Greater tuning engagement is associated with increases in both quantities. Colored curves show population medians. **G**, Motion-dependent modulation across the cumulative model readout for high-tuning-engagement movies. Gain-invariant temporal modulation (left) and SSI sharpening (right) increase as successive feature stages contribute to the output. In **B, F**, and **G**, gray boxes show across-unit interquartile ranges and medians; whiskers show the 5th–95th percentiles across units. Colored points and capped bars show population estimates and 95% bootstrap confidence intervals. Stars indicate positive effects relative to zero improvement (*p <* 0.05, two-sided bootstrap tests with Holm correction across all 32 displayed effects; Methods).

In the illustrative example in Fig. 4A, the moving image produced both a higher firing rate and a more concentrated activation map than the same image held stationary on the retina. Across the population, firing rate and SSI generally increased with the amount of image motion in the input history (quantified by the path length of the eye’s position over the 250 ms history; Fig. 4B). In the sixth of eight equal-count bins over path length (median 22.0 arcmin), measured motion increased firing rate by 23.2% (95% CI [17.5, 29.7]) and SSI by 9.0% (95% CI [5.4, 13.2]) relative to stabilization. Both population effects were positive on average in all eight path-length bins (Holm-adjusted *p <* 0.05; Methods), despite variation in the direction and magnitude of individual unit effects. Thus, the model predicts that not only do FEMs drive additional spikes, they also increase the spatial information carried by each spike.

What property of retinal-image motion produces this effect? Work by Rucci and colleagues has demonstrated that fixational motion creates temporal modulations from spatial image structure [6, 32]. When an image is stationary on the retina, its power lies at zero temporal frequency. Retinal translation redistributes that power into nonzero temporal frequencies determined jointly by the image’s spatial structure and the statistics of motion. Retinal spectra calculated from our measured eye trajectories showed this redistribution for both drift windows with no detected saccade and windows containing microsaccades (Fig. 4C). For the same natural images, microsaccade-containing windows shifted normalized power toward higher temporal frequencies relative to drift. These observations motivated us to test whether the spatiotemporal distribution of retinal power predicts changes in firing rate and spatial information per spike beyond eye-movement path length.

We therefore measured the joint spatiotemporal-frequency (SF × TF) tuning of the pseudo-population model units using simulated drifting gratings (Methods). The units showed a broad range of tuning (Fig. 4E). The contours in Fig. 4C mark the spatial and temporal frequency ranges that strongly drive two example units with contrasting tuning, allowing us to compare how much retinal power falls within each unit’s preferred range. Compared to windows containing only drift, microsaccade-containing windows redistributed normalized natural-image power toward higher temporal frequencies and lower spatial frequencies, increasing the power fraction within one unit’s preferred frequency range while decreasing it within the other’s (Fig. 4D). We therefore expect changes in fixation dynamics to affect units differently depending on their spatiotemporal tuning.

We next asked whether fixation histories that generated more retinal power at a unit’s preferred frequencies also produced larger increases in firing rate and spatial information per spike. For each of the 725 model units, we quantified this overlap by weighting the motion-induced retinal power by the unit’s spatiotemporal tuning and orientation sensitivity. We termed this quantity “tuning engagement” and summarized it across images for each fixation history (Methods). Both firing-rate and SSI modulation increased with tuning engagement (Fig. 4F): the median unit effect rose from 1.7% in the lowest engagement quintile to 63.9% in the highest for firing rate, and from 0.4% to 16.7% for SSI. Within-unit Spearman correlations were larger for tuning engagement than for path length (rate: median *ρ* = 0.891 versus 0.822, paired difference 0.068, 95% CI [0.066, 0.069]; SSI: 0.726 versus 0.690, paired difference 0.044, 95% CI [0.037, 0.047]). These descriptive correlations motivated us to test whether tuning engagement improved prediction of firing-rate and SSI modulation beyond eye-movement path length and the presence of a microsaccade.

We therefore fitted separate regression models for each unit to predict firing-rate and SSI modulation from eye-movement path length and the presence of a microsaccade. These models allowed nonlinear relationships between path length and response modulation that could differ between drift-only and microsaccade-containing windows (Methods). Adding tuning engagement reduced held-out squared prediction error by a median 15.3% for firing-rate modulation (95% CI [6.3, 24.1]%) and 15.5% for SSI modulation (95% CI [10.6, 20.3]%; source-trial bootstrap). These results support a framework in which FEM-induced response enhancement and spatial sharpening depend on how motion-generated retinal power overlaps neuronal tuning. This framework predicts both effects better than movement distance and microsaccade presence alone.

Finally, we noted that increased firing rate is an expected consequence of redistributing retinal power toward frequencies that drive a neuron. It was less clear, however, how this redistribution would increase spatial information per spike. Spectral power omits Fourier phase, whereas the model performs a nonlinear computation on the complete spatiotemporal movie. We therefore considered whether nonlinear processing of the retinal inputs contributed to spatial sharpening. To examine this possibility, we replayed matched moving and stabilized movies through the fitted model and compared predictions formed from first-stage features alone, first- and second-stage features together, and features from all three stages. We measured both temporal modulation, normalized by mean firing rate, and spatial information per spike. The motion-induced increases in both quantities were small for the first-stage contribution and larger when contributions from the second and third stages were included (Fig. 4G). These results suggest that progressive nonlinear processing stages contribute to spatial sharpening by transforming FEM-generated temporal modulation into a more spatially informative response.

## Discussion

Neural activity supporting central vision is continuously shaped by self-generated motion of the eyes, even when gaze is tightly regulated in a task that requires no sampling movement. Measuring this required tracking gaze at the scale of foveal receptive fields while recording dense populations of neurons. Decomposing the population covariance showed that fixational eye movements were the dominant source of rate variability in the foveal population, contributing more than the repeated sequence of natural images we presented.

Conventional assays of neuronal noise (i.e., residual variability) combine internally-generated fluctuations with effects of differences in retinal input that are not captured by the trial-averaged response [16, 17]. In the fovea of alert primates, that treatment misattributes a large fraction of the residual variability, because gaze follows a different trajectory on every trial. Accounting for those trajectories lowered the population Fano factor from 1.25 to a sub-Poisson 0.81 and reduced the mean pairwise correlation from 0.076 to 0.010, consistent with the empirical summaries above. Fixational eye movements accounted for nearly all of the shared variability that the trial-averaged response did not explain.

FEM-driven variability overlapped the dominant dimensions of the stimulus-locked population response, showing that even small eye movements substantially reshape activity within these dimensions across stimulus presentations. Primates nevertheless resolve fine spatial detail during fixation, with psychophysical evidence showing that retinal motion supports spatial resolution [3, 5]. Taken together, these observations motivate a single-trial account of population coding and readout in V1. Variability around the trial average includes systematic responses to image features moving across receptive fields, even when the displayed scene remains unchanged. These responses are part of the visually driven signal, although conventional analyses often assign them to noise.

Measured spike-count correlations span two orders of magnitude across studies, and much of that spread has been attributed to features of the preparation rather than of the circuit, including anesthetic state [33], slow fluctuations in gain [34], firing rate [35], and the counting window and spike-sorting criteria [27]. This study emphasizes that correlations measured during fixation can also reflect shared changes in retinal input due to the FEMs. Our results show that trial-to-trial differences in eye position can inflate estimates of noise correlation in experiments with awake animals. Fortunately, eye position differs from other factors by being readily and directly monitored non-invasively with instrumentation that is now widely available [23–25], and its influence can be accounted for with the decomposition presented here.

We compared the predictive contributions of retinal and explicit eye-state inputs using an image-computable model that predicted individual trials better than the cross-trial average. Removing explicit eye-position and eye-velocity signals left single-trial prediction and eye-driven modulation largely intact, whereas stabilizing the retinal input substantially reduced both. Preserving current retinal position while removing motion within each input history recovered much of the global-stabilization loss. This is consistent with the flashed-image paradigm used to drive neurons: the 3° images changed at 20 Hz, providing temporal modulation even with fixed gaze, with eye position determining which features fell within the small foveal receptive fields (or the sub-fields of simple-type receptive fields). Recent retinal motion made an additional contribution to single-trial prediction, although the impact of removing it was not demonstrably larger than that of the explicit extraretinal input. This supports a substantial reafferent contribution: much of the measured modulation can be predicted from changes in the visual input itself, as suggested by earlier physiological work. For example, Kagan and colleagues showed that retinal image motion and position account for distinct V1 responses to saccades and drifts, alongside extraretinal modulation in around a third of the analyzed neurons [12, 13]. Our measurements and analyses extend this foundational notion of eye movements impacting receptive field drive by quantifying how strongly retinal input predicts single-trial responses, how it accounts for FEM-dependent rate modulation and covariance, and how it affects population-level coding.

This evidence of a major contribution of fine-scale reafference links the variability associated with FEMs to a well-established function of retinal motion. Fixational motion converts spatial structure into temporal modulation, redistributing natural-image power into the temporally band-pass regime of early visual neurons [32, 36], and retinal stabilization impairs fine spatial vision [3, 5]. Our results connect this input transformation to the cortical population code. Measured fixation trajectories produced both stronger responses and greater spatial information per predicted spike than when the model was stimulated with perfectly stable inputs. Considering the engagement based on spatiotemporal tuning added predictive information about both effects beyond movement distance and microsaccade classification. After controlling total dynamic power, normalized spectral overlap retained a modest predictive benefit for SSI. This supports a contribution of the retinal signal’s spectral distribution to model spatial sharpening beyond these movement descriptors and the overall amount of modulation. It does not establish unique neuron–tuning routing, or exclude other descriptions of eye-movement dynamics.

The space-to-time spectral redistribution describes the signal generated by FEMs, but not the full computation that produces a sharper spatial representation. The spatiotemporal tuning analysis retains power but discards Fourier phase, whereas the fitted model was directly image-computable on the spatiotemporal movie. Adding successive feature-stage contributions to the trained readout increased both temporal modulation and spatial information for high-tuning-engagement movies. In the fitted model, FEMs make spatial structure available as a temporally modulated signal, and subsequent transformations produce a more spatially informative response. The specific spatiotemporal tuning and stagewise transformations identified here are model-derived predictions that can be further tested with controlled dynamic stimuli.

A downstream circuit could amplify or attenuate these changes depending on which V1 responses it pools. Attention models make the relevant distinction explicit: modulation aligned with the gradient of task performance can be more effective than modulation based on neuronal tuning alone [37]. Analogously, FEM-dependent changes could have larger effects downstream when they engage inputs to which a later computation is sensitive. This suggests a testable connection to attention: changes in fixation dynamics may alter the retinal signals available to downstream selection. Because our dataset did not involve manipulating attention or measuring downstream responses, our results only lay the conceptual and technical groundwork for additional work to consider FEMs in the context of effects typically attributed completely to attention.

Active vision is often considered at the scale of saccades, with each fixation treated as a static sample [38–41]. Our results instead show that active sensing continues throughout fixation. Movements of fractions of a degree account for much of the apparent variability in foveal V1 while simultaneously generating signals that improve spatial encoding. Accounting for retinal image motion substantially reduces how much cortical variability should be assigned as unexplainable noise, and reveals a visual signalling computation that sharpens spatial representations.

## Methods

### Animals and surgical preparation

Two adult male common marmosets (*Callithrix jacchus*), monkeys L and A, contributed the recordings analyzed here. The monkey L recordings were collected as part of the dataset described by Yates et al. [23]. The monkey A recordings were collected subsequently on the same preparation and acquisition system, with the modifications noted below. The fixated flashed-image condition was not part of that report.

All surgical and experimental procedures were approved by the Institutional Animal Care and Use Committee at the University of Rochester and conformed to US National Institutes of Health guidelines. Each animal was implanted with a titanium head post and a recording chamber positioned over the foveal representation on the lateral surface of V1 [23].

### Eye tracking and stimulus presentation

Animals were head-fixed during recording and viewed the display through a hot mirror. Eye position was measured with a digital dual Purkinje image eye tracker [23–25]. Visual stimuli were generated in MATLAB using Psychtoolbox [42] and presented with a ProPixx projector (VPixx Technologies). Stimulus and electrophysiology clocks were synchronized with a Datapixx interface, which transmitted event words to the Open Ephys acquisition system. Stimulus code is available at https://github.com/jcbyts/MarmoV5.

### Fixated flashed-image condition

Animals maintained fixation while viewing a repeated sequence of whitened natural images. Images subtended 3° of visual angle and were presented at 20 Hz. Analyses were restricted to periods in which valid eye-position estimates remained within 0.5° of the fixation point, a 1°-diameter fixation window. Repeated presentations of the same sequence provided matched stimulus epochs with naturally varying fixational eye trajectories.

### Free-viewing stimulus conditions

Each session also included three free-viewing conditions taken from the foraging paradigm described previously [23]. No fixation was required in these conditions. The animal viewed the display freely while stimuli were presented continuously, and small Gabor or face targets appeared intermittently at varying screen locations to encourage saccades. Eye position was tracked throughout, and the stimulus falling on each receptive field was reconstructed offline in retinal coordinates. These conditions were not used for the variability decomposition. They provided the receptive-field estimates used to select units for analysis, the training data for the encoding model, and the natural free-viewing fixations analyzed in Figure 4.

#### Gabor noise

A dense field of Gabor patches covered the display, and a new set of patches was drawn on each noise frame, with the orientation, phase, spatial frequency, size, and position of every patch drawn independently at random. This condition drove units across a wide range of orientations and spatial frequencies, and provided the data used to compute spike-triggered averages and spike-triggered energies.

#### Gratings

A full-field sinusoidal grating was presented at fixed phase and contrast, with its orientation and spatial frequency redrawn on each noise frame.

#### Natural images

A single natural image spanning the display was presented on each trial and viewed freely. Fixations extracted from this condition provided the eye-position distributions and local image measurements analyzed in Figure 4.

### Electrophysiology and spike sorting

Neural activity was recorded with multisite silicon electrode arrays (NeuroNexus). Each array contained two shanks separated laterally by 200 *µ*m, with 32 recording sites per shank at 35 *µ*m spacing. Arrays used for monkey A were coated with poly(3,4-ethylenedioxythiophene) (PEDOT) to reduce impedance and improve signal quality [43]. Wideband signals were amplified and digitized at 30 kHz with Intan headstages using Open Ephys.

Recordings from monkey L were processed and spike sorted with Kilosort 2 following Yates et al. [23], including manual curation in Phy. Recordings from monkey A were processed with an updated pipeline and spike sorted with Kilosort 4 [44]. Both single-unit and multi-unit clusters were considered for analysis.

Units were included if they exhibited a visually driven receptive field, defined by more than 100 spikes during receptive-field mapping and a peak spike-triggered energy SNR greater than 5. That SNR was the maximum spatial deviation of the smoothed energy map from its median, normalized by the median deviation at the earliest lag.

### Decomposition of neural variability

Trial-to-trial variability in neural responses reflects stimulus-locked structure, variability driven by eye movements, and residual variability that remains once both are accounted for. We separate these components by applying the law of total covariance twice to the measured spike-count covariance [15].

#### Setup and notation

Let 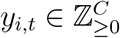 denote the spike counts of the *C* simultaneously recorded units on trial *i* at stimulus time *t*, and let *e*_*i,t*_ denote the eye position on the same trial at the same time. Every trial repeats the same frozen stimulus sequence, so *t* indexes the same stimulus frame across repeats. The dimension of *e*_*i,t*_ is left unspecified, as it may represent eye position alone or a short trajectory history, and could in principle contain other covariates such as a second eye variable, velocity, or other potentially relevant behavioral variables.

We write *Y* for *y*_*i,t*_ treated as a random vector over trials and times. Subscripts on E and Cov name the variables an operator ranges over, so Cov_*i,t*_(*Y*) is taken across both trials and stimulus times, while Cov_*i*_(*Y* | *t, e*) is taken across trials at a fixed stimulus time and eye state. Each covariance is a *C × C* matrix over units whose (*c, d*) entry is the covariance of the spike counts of units *c* and *d*. The total population covariance is the covariance of spike counts across trials and time,

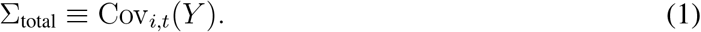

#### Separating residual and rate covariance

Applying the law of total covariance while conditioning on both stimulus time *t* and eye state *e* yields

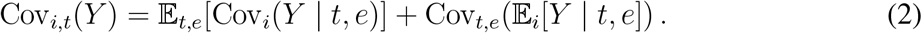

The first term averages the trial-to-trial covariance of spike counts at fixed *t* and *e* over all times and eye states. It is the variability that remains after conditioning on every measured external variable, which we call the residual covariance,

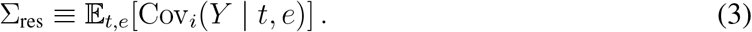

The second term is the covariance of the conditional mean spike count, *R*(*t, e*) ≡E_*i*_[*Y* | *t, e*], which we call the rate covariance,

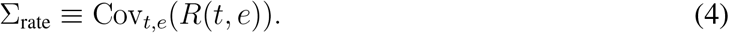

The total covariance therefore separates into a residual component and a component arising from structured changes in the underlying firing rates [45],

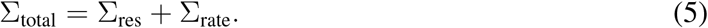

#### Separating eye-movement and stimulus-locked rate covariance

Applying the law of total covariance a second time, now to the rate term *R*(*t, e*) and conditioning only on stimulus time *t*, yields

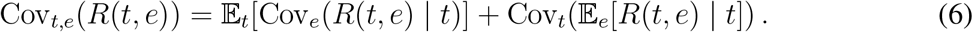

The first term is the variability in firing rate at a fixed stimulus frame that arises from trial-to-trial differences in eye state, which we call the fixational eye movement (FEM) covariance,

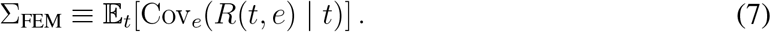

The second term depends only on the stimulus-locked mean rate,

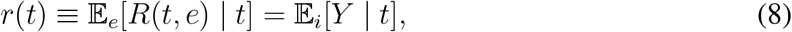

and is therefore driven by the frozen stimulus sequence alone. We call it the PSTH covariance,

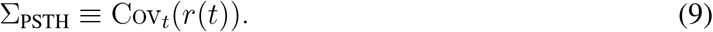

The rate covariance therefore separates into an eye-movement component and a stimulus-locked component,

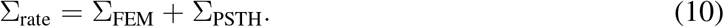

#### Final decomposition

Combining the two applications of the law of total covariance yields the full factorization

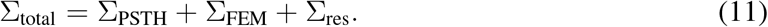

The population covariance is the sum of three components: covariance driven by the stimulus-locked response, additional rate covariance induced by fixational eye movements, and residual covariance. The identity requires only that spike counts have finite second moments, and makes no parametric assumption about how firing rate depends on eye position. The further assumptions needed to estimate the three terms are given with the estimators below.

Conventional analyses define residual covariance relative to the trial-averaged response [28, 46],

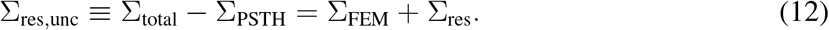

We call this quantity the uncorrected residual and Σ_res_ the residual.

### Estimating the covariance components

We estimated Σ_PSTH_ and Σ_rate_ from a common principle. For two distinct trials *i* ≠ *j* at the same stimulus time *t*, the residual noise is independent, so the product of their spike counts is an unbiased estimate of the product of their underlying firing rates, with the observation noise cancelling in expectation. The two terms differ only in which pairs enter the average, all same-time pairs for the stimulus-locked term and eye-position-matched close pairs for the rate term. Estimating both from the same distinct-trial products, with a shared time-bin weighting and a shared mean, lets the four terms compose exactly and treats non-stationary drift in firing rate identically across terms.

Two features of the data can bias these estimators. The first is a correlation between image content and eye position. Because eye position is distributed around a central tendency at the middle of the fixation window, trajectories are more likely to be similar near the center of that distribution, so for a single image the trajectory distance between two trials is correlated with the image features at that position. We protect against this by presenting many images rapidly, so that no feature is consistently paired with the center of the fixation window. The second is slow drift in firing rate, which changes the sample mean across time for reasons unrelated to the stimulus or eye movements. We estimated Σ_PSTH_ and Σ_rate_ against a single global mean, computed under the same pair-count time-bin weighting and shared by both terms, so that drift contributes identically to both and cancels in their difference, 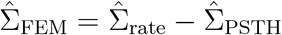.

Such drift is nonetheless consequential for the correlations we report. Because close pairs are matched on eye trajectory rather than on proximity in recording time, shared slow fluctuations survive into 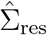 as well as 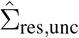, so the FEM-corrected correlations are an upper bound and the near-zero values we obtain are conservative.

#### Consistent time-bin weighting

Fixation durations varied across trials, so after aligning to fixation onset the number of contributing trials *n*_*t*_ decreased across stimulus time bins. The three terms have different natural weightings over these bins: the close-pair rate estimator weights each bin by its pair count (∝*n*_*t*_(*n*_*t*_ *−* 1)/2), a per-bin-average PSTH weights each bin equally, and the pooled sample covariance weights each bin by *n*_*t*_. When *n*_*t*_ varies these weightings disagree, and the resulting bias falls on the off-diagonal (cross-unit) covariance, distorting the estimated noise correlations while leaving the per-unit variance almost unchanged. We therefore pinned every term, including the mean subtracted from each, to a single pair-count weighting, so that the law of total covariance holds term by term.

#### Eye-trajectory shuffle

We evaluated all analyses against surrogate data in which the link between eye movements and spikes was broken by permuting eye trajectories across samples, where a sample is one spike-count window from one trial at one stimulus time. The permutation breaks the pairing across both trial and time, while spike counts, 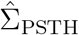, and the per-unit mean rates are held at their observed values. We drew 1,000 surrogates per session and recomputed 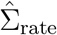, and every quantity derived from it, on each. For Fano factor, noise correlation, and subspace alignment (Fig. 2E,F,I), empirical *p*-values compare the observed across-session statistic with the corresponding surrogate distribution, using a plus-one finite-sample correction. The FEM-fraction distribution (Fig. 2C) is summarized by its pooled unit median and a 5,000-resample unit-bootstrap confidence interval; no pooled-population shuffle significance test is reported for that panel.

#### Total covariance

We estimated Σ_total_ as the sample covariance of the raw spike counts across all trials *i* and stimulus time bins *t*, under the common time-bin weighting above,

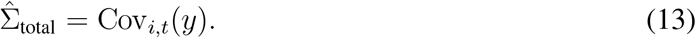

This term captures all sources of variance, including time-varying mean rates, eye-movement modulations, and Poisson-like noise. Because it pools across time and trials, it is well estimated even for units with low firing rates.

#### Stimulus-locked covariance

The sample covariance of the peri-stimulus time histogram (PSTH) is a biased estimate of the stimulus-locked covariance, because each trial-averaged mean still carries the sampling noise of a finite trial count. We removed this bias with the distinct-trial product introduced above [47]. At each stimulus time bin *t* we averaged the outer product of spike counts over all pairs of distinct trials *i* ≠ *j*, averaged the result across time bins, and subtracted the outer product of the global mean rate 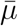, computed across all trials and time bins under the common weighting rather than per bin,

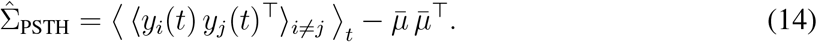

Because *i* and *j* are different trials, their residual noise is independent and averages away, leaving an unbiased estimate of the covariance of the trial-mean rate across stimulus time.

#### Eye-movement-dependent covariance

We estimated Σ_FEM_ indirectly, by first estimating the total rate covariance Σ_rate_, which captures all variance due to time-varying firing rates, both stimulus-driven and eye-movement-driven, and then isolating the eye-movement component by subtraction,

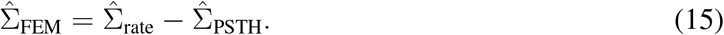

To estimate Σ_rate_ we again averaged distinct-trial products, but restricted the average to pairs whose eye trajectories nearly coincided. If two trials *i* and *j* follow the same eye trajectory up to stimulus time *t*, their rates conditioned on stimulus, time, and eye position are equal, so the product of their spike counts estimates the squared rate with the residual noise removed. For each time bin across pairs of distinct trials we computed the root-mean-square (RMS) distance between eye trajectories over the history window of duration *H*,

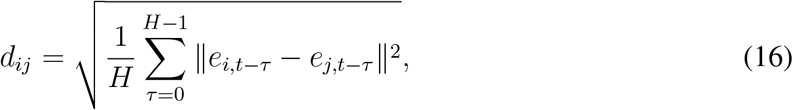

pooled all pairs closer than a threshold *d*_*ij*_ *< ε* (*ε* = 0.05°), and used this pool to compute an estimate of the variance conditioned on there being little difference in the eye’s recent trajectory between trials,

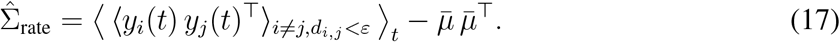

The 0.05° criterion approximated conditioning on identical trajectories. Because *ε* is small but finite, the result is a mildly conservative estimate of the rate covariance at *d*_*ij*_ → 0. As formulated, this is equivalent to the analysis used by [15].

#### Correcting the close-pair sampling bias

Conditioning on close pairs biases Σ_rate_ when the stimulus is not spatially homogeneous. The two eye positions of a close pair are each drawn from the fixational distribution *p*(*e*) and must nearly coincide, so close pairs occur in proportion to *p*(*e*)^2^, a tighter distribution concentrated toward the center of fixation, whereas Σ_PSTH_ and Σ_total_ average over the full viewing distribution *p*(*e*). If firing rate were independent of absolute eye position, as for a spatially stationary stimulus, this would not matter, and the close-pair second moment would recover the same rate variance under either distribution [15]. That condition does not hold here. At each stimulus frame the image is fixed and identical across trials, so a unit’s rate depends on where the eye sits within it, and the close-pair and full distributions yield different rate variances. Combining Σ_rate_ estimated over one with Σ_PSTH_ and Σ_total_ estimated over the other breaks the factorization.

We corrected this by importance-reweighting the close-pair second moment onto the true viewing distribution. We estimated the fixational density 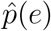 and the density of close-pair midpoints 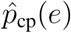 by kernel density estimation, and weighted each close pair at midpoint *e* by the ratio 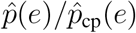, normalized across pairs and clipped to bound the influence of rarely visited positions. This maps the close-pair sampling density back to *p*(*e*), so that Σ_rate_, Σ_PSTH_, and Σ_total_ are all quantified over the distribution the animal actually viewed. The correction extends the cross-trial decomposition, which assumes a spatially homogeneous stimulus, to structured stimuli whose rate variance depends on where the eye lands.

### Neural variability metrics

Metrics were computed for units with mean firing rates above 2 spikes/s and split-half PSTH reliability *R*^2^ *>* 0.10 [31]. We defined *R*^2^ as the squared Pearson correlation between split-half PSTHs, averaged across 100 random trial splits. Sessions with fewer than 10 qualifying units were excluded. These criteria retained a mean of 79.6 ± 26.1 units per session for monkey A and 18.2 ± 9.1 for monkey L (mean ± s.d. across sessions). Primary results used a 25-ms counting window; analyses at 8.3, 16.7, and 50 ms assessed robustness to counting-window duration.

#### Population Fano factor

We quantified population variability from the relation between residual spike-count variance and mean spike count across units within each recording session. For unit *c*, the uncorrected and FEM-corrected residual variances were the corresponding diagonal entries of 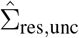 and 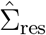, respectively. For each condition, the population Fano factor was the slope through the origin of residual variance *v*_*c*_ against expected spike count *µ*_*c*_ [48],

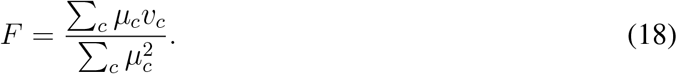

Slopes were computed separately for each session. Figure 2E reports the mean and standard deviation across sessions. Statistical significance was evaluated against the eye-trajectory shuffle, recomputing the FEM-corrected slope within each session and comparing the observed across-session mean reduction with the surrogate distribution.

#### Noise correlations

Pairwise noise correlations were computed separately from 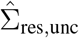 and the FEM-corrected 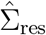. Before conversion to correlations, each covariance matrix was projected onto the positive-semidefinite cone by setting negative eigenvalues to zero. For units *c* and *d*, noise correlation was defined as

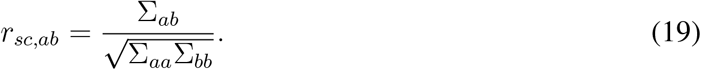

We retained the upper-triangular entries corresponding to unique unit pairs. Correlations were averaged across pairs within each session, and session means were then averaged with equal weight across sessions. Confidence intervals were estimated with 5,000 bootstrap resamples across sessions, and the FEM-corrected minus uncorrected contrast was calculated within each session. Under the eye-trajectory shuffle, corrected noise correlations and their within-session means were recomputed and averaged across sessions, and the empirical test compared the observed across-session mean change with that surrogate distribution.

### Alignment between stimulus-locked and eye-movement covariance

We measured whether Σ_FEM_ occupied the same population dimensions as Σ_PSTH_, using the 25 ms counting window of the variability metrics above. Both matrices were projected onto the positive semi-definite cone as described above and eigendecomposed, 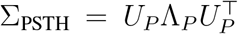 and 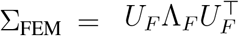, with eigenvalues in descending order. We retained the leading *k* eigenvectors of each, *U*_*P,k*_ and *U*_*F,k*_ ∈ ℝ^*C×k*^, for a population of *C* units.

Alignment was quantified as the fraction of one component’s variance lying in the other’s leading subspace,

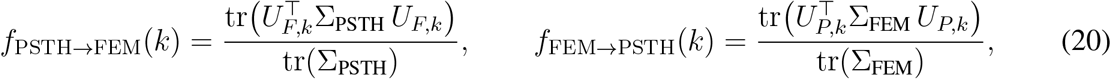

the fraction of stimulus-locked variance captured by the FEM subspace and the fraction of FEM variance captured by the stimulus subspace. The measure is asymmetric, so we report both directions.

We set *k* = 4, matching the effective dimensionality of the two components. That dimensionality was measured by the participation ratio, (tr Σ)^2^/tr(Σ^2^) [49], which averaged 3.6 for Σ_PSTH_ and 3.4 for Σ_FEM_ across sessions (Figure 2G).

Significance was assessed against the eye-trajectory shuffle. Each surrogate recomputed 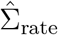, and with it Σ_FEM_ and its eigenvectors, while Σ_PSTH_ and its subspace were held at their observed values. Figure 2I reports the across-session mean ± SD against that surrogate distribution. A null built on random orientation would set a much lower bar. A random *k*-dimensional subspace captures *k/C* of a component’s variance, which averaged 0.14 across sessions, whereas under the eye-trajectory shuffle the same measures averaged 0.35. Testing against random orientation would therefore overstate the significance of the observed alignment.

### Encoding model

#### Model architecture

The model follows a “core-readout” architecture [50] with a shared feedforward visual core, a separate extraretinal/behavioral pathway, and session-specific readouts. The fitted rank-one model contains 5,295,486 parameters: 3,421,516 in the visual core, 46,520 in the extraretinal/behavioral pathway, and 1,827,450 in the readouts. It predicts 2,790 recorded output channels across 30 sessions; the unit subsets entering each figure are specified separately.

#### Retinal pathway

Gaze-corrected grayscale stimuli [23] were center-cropped to 35 × 35 pixels and retained at their native 240-Hz sampling rate. Each prediction receives 60 frames of causal stimulus history (250 ms). Fourteen learned 60 × 7 × 7 spatiotemporal filters collapse this history into a single feature map. Three subsequent spatial convolutions use 84 signed channels each and kernels of 15 × 15, 11 × 11, and 9 × 9 pixels. Group normalization and local response normalization precede a sign split and rectification, yielding 168 nonnegative channels per spatial stage. Spatial kernels are Hamming-windowed; the first-layer filters additionally use a Hann frequency mask in time and space. Between spatial stages, padded 3 × 3 pooling with stride 2 and an *L*^5^ norm reduces spatial resolution. Nearest-neighbor resizing brings the three stage outputs to a common 9 × 9 grid, and their concatenation provides a 504-channel multiscale representation.

#### Extraretinal pathway

The behavioral input contains 42 features: horizontal and vertical eye position plus 40 sign-split velocity features. Position differences are normalized, transformed with a symmetric logarithm, and projected onto ten raised-cosine basis functions per axis, with peaks spanning 30–200 ms. The bases are acausal, giving this pathway access to eye velocity preceding and following the prediction time. A multilayer perceptron with 128 hidden units and GELU nonlinearities embeds the behavioral vector into 64 features. A bounded gain, 1 + 0.5 tanh(*g*), modulates each multiscale visual channel; the 64 behavioral features are also broadcast across space and appended as additive readout channels. The model ablations manipulate this pathway and the gaze-contingent retinal input independently.

#### Sparse factorized readouts

Each session has separate readouts for its recorded units, initialized from Gaussian spatial masks [51] and subsequently optimized as sparse spatial and feature factors. The readout receives all 504 multiscale visual channels and 64 appended behavioral channels on the 9 × 9 grid. The readout has rank *R* = 1. For neuron *i*, the predicted count is

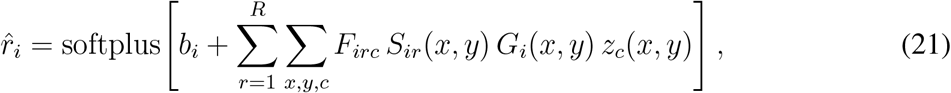

where *z*_*c*_ is a modulated visual or appended behavioral feature, *F*_*irc*_ is a feature weight, *S*_*ir*_ is a learned spatial factor, and *G*_*i*_ is a learned Gaussian envelope shared across the neuron’s factors. Each neuron therefore uses one spatial–feature product. The Gaussian center, widths, and orientation are learned. A single per-neuron bias and the softplus link produce the expected spike count in each 1/240-s bin.

#### Training data and evaluation protocol

The model was fit to Gabor noise, gratings, and free-viewing natural images pooled across 30 recording sessions from the two animals. The initial trial split allocated 70% to training, 15% to validation, and 15% to testing. Natural image stimulus splits remained fixed throughout fitting. During the second and third training stages, all genuine grating repeats were exposed to training and gratings were sampled with weight 16 relative to the other stimulus types. Recorded-grating capture by the final model is therefore descriptive and is not a held-out generalization result.

The fixated flashed-image condition was withheld entirely from model fitting. Figure 3 evaluates predictive accuracy on repeated natural images flashed at 20 Hz under enforced fixation, a stimulus and behavioral context absent from training. All trials passing the evaluation filters are eligible because this condition contributed no training data. Figure 4 instead uses controlled drifting-grating assays and natural-image replays through the fitted model; these counterfactual analyses characterize the model and are not additional held-out neuronal prediction tests.

#### Training procedure

We minimized the Poisson negative log-likelihood, evaluated only at time points passing the data filters,

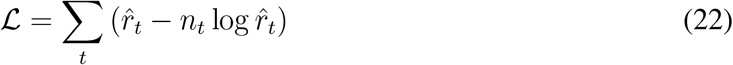

where 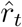 is the expected count per native time bin and *n*_*t*_ is the observed spike count. Optimization used AdamW, a two-epoch linear warmup followed by cosine annealing, bfloat16 mixed precision, and gradient-norm clipping at 10. Batches contained 128 samples from a single session, with sessions sampled uniformly, and an epoch comprised 512 batches. Weight decay was 10^−5^.

The model was fitted in three training phases using seed 201. First, the visual core, behavioral pathway, and Gaussian readouts were trained from scratch for 488 epochs at an initial learning rate of 5 × 10^−4^; the checkpoint with the best validation bits per spike is retained. Second, the shared visual and behavioral representation is frozen, and the Gaussian heads are embedded in sparse rank-one readouts while preserving their predictions at the handoff. Only the readouts are optimized for 24 epochs at 10^−4^, and the final checkpoint is retained to complete the prescribed sparsity schedule. Third, all components are unfrozen for 64 epochs, with a readout learning rate of 10^−4^ and a core learning-rate multiplier of 0.05; the best validation checkpoint is selected. Every phase uses the same Poisson objective. A weak first-layer spatial Laplacian penalty encourages smoothness.

Proximal L1 sparsity is applied only to the readout’s feature and spatial weights during the second and third training phases. The feature and spatial penalty coefficients are 0.1 and 0.02, respectively. Their strengths ramp from zero to their full values over epochs 2–12 of the second phase and remain constant in the third phase. After each optimizer update, each neuron’s spatial factors are normalized jointly over rank and spatial position before scalar-learning-rate L1 soft thresholding is applied to the spatial and feature weights. The learned Gaussian envelope multiplies the spatial factors; its widths are constrained to 0.75–2.0 grid pixels during these sparse-readout phases. The first training phase does not apply proximal L1.

#### Model ablations

We evaluated three conditions using the same fitted model weights. The full condition retained both the gaze-contingent retinal stimulus and the extraretinal eye-position and eye-velocity inputs. The retinal-only condition set the extraretinal inputs to zero while retaining the gaze-contingent retinal sequence. The stabilized-retina condition retained the extraretinal inputs but froze the retinal crop for every trial within a session at one common eye position. We defined this position as the centroid of valid eye-position samples within 0.5° of fixation and used the recorded eye-position bin nearest that centroid to render the fixed crop. Using one session-global location prevents trial identity from remaining in the retinal input through trial-specific stabilization points.

We additionally tested history-local stabilization (Extended Data Figure 4). For each 240-Hz-native prediction, we rendered all 60 input frames at the ROI of that prediction’s latest input sample, retaining each historical frame’s actual flashed image and display position. Thus the current retinal frame is identical to the measured input, while preceding gaze displacement is removed. This is a history-specific counterfactual, not one globally consistent stabilized movie: retinal position can change from one prediction to the next. Both stabilization controls retain the measured behavioral inputs and the temporal modulation from the 20-Hz image sequence. Rendering was checked against the native renderer and every current frame was verified pixel-for-pixel. We used the same canonical observations, validity masks, neuron identities, correlation ceilings, affine calibration procedure, and conditional-rate-variance denominators as Figure 3C,D. Full and retinal-only inputs were replayed as numerical reproducibility controls; the additional analysis did not recompute the FEM variance decomposition. To compare ablation costs, we subtracted each ablated score from Full within each unit. The difference between the recent-motion and extraretinal costs is therefore the Retinal-only score minus the History-local score; we summarized this paired contrast directly, rather than subtracting population medians. Paired effects were summarized with 10,000 session bootstrap resamples, conditional on the fitted model.

### Model performance metrics

We evaluated predictive performance on the withheld fixated flashed-image condition with three metrics. Noise-normalized correlation (CC_norm_) measures trial-averaged accuracy, the captured fraction of conditional rate variance quantifies trial-by-trial predictability, and the FEM modulation fraction tests whether the model reproduces the empirical eye-movement dependence. Metrics were computed per unit on matched valid bins and summarized across the eligible units and sessions.

#### Noise-normalized correlation

To benchmark stimulus-locked predictive accuracy with a standard metric, we report the noise-normalized correlation coefficient CC_norm_, matching the definition used by [50] to enable direct comparison. Let *r* denote the in silico prediction and *y* the in vivo response, and let ⟨·⟩ denote averaging across stimulus repeats (trials). Define the Pearson correlation between mean responses

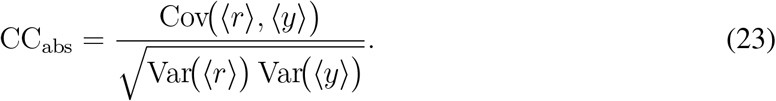

Following [31], an upper bound on achievable correlation given finite trial count *N* is

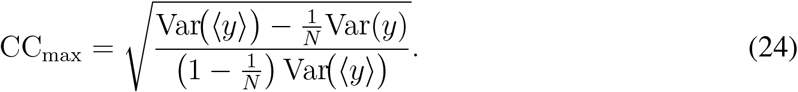

We then report

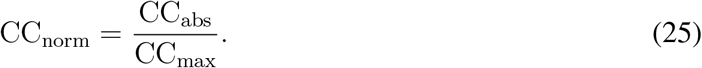

#### Conditional rate variance captured on single trials

For each unit and predictor, we quantified captured count variance as

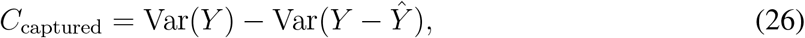

where *Y* is the observed spike count and *Ŷ* is either a model prediction or the leave-one-out trial-average prediction. The model predicts at 240 Hz. Native observed and predicted counts were summed in endpoint-aligned pairs to match the fixed 120-Hz empirical analysis grid, preserving trial boundaries and exact unit identities. We then used one-bin (8.33 ms) counting windows from the Figure 2 analysis within 0.5° of fixation, each carrying the same fixed three-bin (25 ms) matching history that the Figure 2 decomposition applies at every counting window. This counting window is the empirical comparison resolution, not the model’s native bin width. All predictors were scored on the same bins with valid responses, data filters, and predictions, and units with fewer than 50 scored windows were excluded. Before scoring, each condition’s predictions were given a per-unit affine calibration, a gain and an offset fit by Poisson likelihood on the same bins. This places the conditions on a comparable count scale at the cost of in-sample optimism, applied identically to every condition.

We normalized captured variance by the unit’s conditional rate variance from the Figure 2 covariance decomposition,

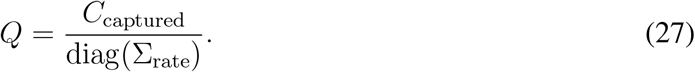

Writing the observed count as *Y* = *λ* + *ε*, where *λ* = E[*Y* | stimulus, eye position] is the conditional rate and *ε* is the conditional residual, the residual is uncorrelated with any function of the conditioning variables. Under matched conditioning and sampling, *Q* therefore reduces to

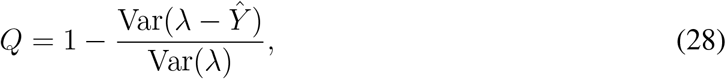

an ordinary variance-explained coefficient whose target is the conditional rate rather than the noisy spike count. Any constant prediction scores zero, and a prediction that matches *λ* up to an additive constant scores one. Because the denominator is fixed across conditions within a unit, normalization cannot reorder the conditions; it places units with different noise levels on a common scale before pooling. The denominator is an estimate from the Figure 2 decomposition on its empirical support, whereas the numerator additionally requires valid model predictions. Consequently, finite-sample estimates can fall below zero or exceed one; these values were retained rather than clipped.

Panel-D contrasts were tested with a session-clustered bootstrap. Sessions were resampled with replacement 10,000 times, the units of the resampled sessions were pooled, and the median paired difference was recomputed, giving a percentile confidence interval. Sessions rather than units were resampled because units recorded together share a stimulus sequence and an eye trace. An equal-session-weight sensitivity analysis also gave intervals excluding zero for the three reported contrasts. The full model exceeded the trial average in 16 of 19 session medians, and retinal stabilization reduced performance in all 19 sessions.

#### Model FEM modulation fraction

We computed the FEM modulation fraction for observed and model responses using the same matched close-pair estimator and fixation frame used in Figure 2,

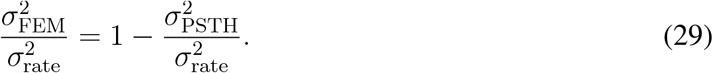

The estimator was applied separately to the full, retinal-only, and stabilized model rates after alignment to the empirical analysis grid. It was run at the 25 ms counting window Figure 2 reports, with the same three-bin matching history and close-pair criterion on RMS trajectory distance. The empirical distribution therefore measures the same quantity as Figure 2C, restricted to the eligible Figure 3 population. Single-trial captured variance above instead uses an 8.33-ms window; neither analysis changes the model’s native 240-Hz prediction rate. Distribution summaries retained finite fractions within [0, 1] without clipping. Equivalence was evaluated separately for each model condition on units with both empirical and model fractions in this range, using paired two-one-sided *t* tests on the mean empirical-minus-model difference with bounds [−0.10, 0.10]. The reported equivalence *p*-value is the larger of the two one-sided values; *p <* 0.05 establishes equivalence within this margin. The stabilized comparison additionally used a paired two-sided Wilcoxon signed-rank test. These unit-level tests do not resample recording sessions; the shaded median band in Fig. 3E is a visual reference rather than the test statistic.

### Spatial information in model-derived populations

For each of the 725 fitted unit readouts, we replicated its spatial pooling over retinal position to form a population of identically tuned model units differing only in position. Each natural image patch was viewed along a measured eye trajectory and paired with a stabilized counterfactual. Recorded-unit identities were retained throughout.

### Retinal-motion replay and spatial information analysis

#### Fixational eye-movement histories

Fixational eye-movement histories were extracted from the continuous free-viewing eye traces [23]. Eye position was placed on a uniform raw-sampling grid and filtered continuously with a symmetric positive Gaussian kernel of standard deviation 6 ms, truncated at five standard deviations. Filtering used 0.6 s of continuous padding on each side before sampling at 240 Hz. This gentle rolloff (approximately 22 Hz at −3 dB) was chosen near the independently measured spectral breakpoint. A positive kernel avoids introducing overshoot around rapid eye movements. Retinal-power spectra were also evaluated with standard deviations of 4 and 8 ms as sensitivity controls. The candidate bank of 1,000 fixation histories sampled across all 30 recording sessions and across the observed path-length distribution. Saccade times and sample indices were aligned to the continuous eye tracking records, preserving sample indices when invalid samples were excluded. Eligible windows required valid native tracking, no unexplained rapid movement, and coverage inside the interval spanned by verified saved events. Native invalid samples and gaps were guarded by 100 ms; event boundaries were guarded by 50 ms. Histories with events of amplitude at least 1°, or with net displacement at least 1° within 50 ms, were excluded from the analysis. Each retained analysis interval contained 60 samples (250 ms). Eye position was centered within the analysis interval. The filtered path length was

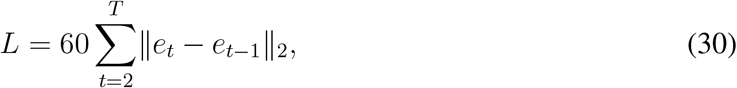

in units of arcmin. An independent screen flagged unlabeled rapid motion when native eye speed after 6-ms Gaussian smoothing exceeded the larger of 5 deg/s and the session median plus 21 median absolute deviations, outside verified event intervals expanded by 50 ms. Flagged histories were excluded rather than assigned a new microsaccade label. The population response analyses pooled eligible drift and microsaccade histories; the illustrative panel-A selection favored verified microsaccade windows. Panels 4C and 4D explicitly compared the two event-defined conditions.

#### Natural-image replay population

The replay population comprised the 725 units with exact session and cluster matches between the analysis population and the fitted checkpoint. Units were retained one-to-one using their exact session and cluster identities.

For the population response matrix, 40 natural-image patches were selected at evenly spaced positions from an independently ordered bank of 100 patches. Two hundred eligible fixation histories were selected after imposing a 350-arcmin maximum path length, with 100 histories from each animal and evenly spaced source rows within animal. Crossing images and trajectories produced 8,000 measured-motion movies. For each natural image, a matched stabilized movie held retinal displacement at zero. The complete behavioral input was fixed to zero in both conditions.

Each movie contained 60 scored samples at 240 Hz. The unavailable causal history preceding the first sample was filled by holding the initial gaze position fixed. The model was evaluated convolutionally over a larger retinal field, allowing each fitted neural readout to be translated over all admissible spatial positions and thereby producing a two-dimensional activation map at every time point. Model outputs are expected spike counts per 1/240-s bin. We multiplied them by 240 to express firing rates in spikes/s and summed the bin counts to obtain the expected count over each 250-ms movie.

The illustrative example in Figure 4A was selected from 12 images with the largest central image-gradient RMS, measured in the central half of the rendered field before model evaluation, using the rate, SSI, and response-map criteria specified below and was not used for population inference. Candidate 250-ms trace windows came from eligible records in the separately filtered one-second fixation bank. Up to six bounded windows per animal were crossed with the 12 candidate images and scored for all 725 units; the final example was restricted to the eight units with the strongest joint ranks of rate and SSI effects in the full population replay. Baseline rate and SSI had to be at least 0.5 spikes/s and 0.05 bits/spike. The strict display gates required rate increases of 25–200%, SSI increases of 10–200% and 0.02–0.25 bits/spike, a moving rate no greater than 50 spikes/s, and normalized map-RMS change of at least 0.12. The final choice maximized the absolute SSI increase within those bounds. We replayed and visually inspected the eight highest-ranked distinct image–unit candidates, retaining the highest-ranked example. For this visualization, the measured and stabilized histories were aligned at the frame associated with the model’s mean first-layer peak-energy lag: 15 frames, or 62.5 ms, before the prediction. The two histories therefore shared the retinal frame expected to contribute most strongly to the displayed response while differing in their preceding motion.

#### Single-spike information

Let *r*_*t*_(*x*) denote a unit’s nonnegative predicted firing rate in spikes/s at spatial position *x* and time *t*, and let 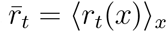. Spatial single-spike information was

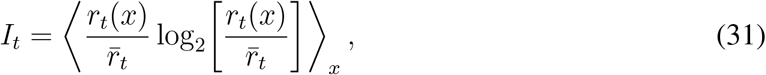

in bits per spike. This quantity is invariant to uniform multiplicative gain and therefore measures spatial response sharpening independently of mean firing-rate gain.

SSI was pooled across time using the expected spike count 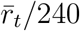 as its weight. Population SSI was likewise pooled across images, trajectories, and units by expected spike count. Rate and SSI effects were reported relative to the matched stabilized condition:

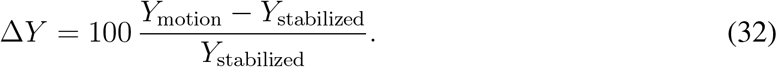

#### Population response versus fixation path length

For Figure 4B, the 200 fixation histories were divided into eight equal-count bins according to filtered path length. All 725 units and all 40 images contributed to every bin. Population firing rate was computed from total expected responses. Population SSI was computed from total spatial information divided by total expected spike count. Stabilized baselines were matched by image and unit.

The colored curves show the spike-weighted population estimand. Confidence intervals were obtained from 3,000 crossed bootstrap resamples of images, fixation histories, and units. The accompanying boxes show the distribution of independently calculated unit effects: the box spans the interquartile range, its center is the unit median, and the whiskers show the 5th and 95th percentiles. Units outside the whiskers remained in all population calculations.

#### Model spatiotemporal tuning

Spatiotemporal tuning was measured by replaying controlled drifting gratings through each unit’s fitted readout at 240 Hz. Each stimulus contained the model’s complete 60-frame history on a 35 × 35-pixel aperture sampled at 37.50 pixels/degree. The bank contained nine spatial frequencies from 1.072 to 15.940 cycles/degree; 14 nonzero temporal frequencies from 1 to 90.51 Hz plus a static condition; 18 motion directions spanning 360°; and 24 uniformly spaced endpoint phases. The primary assay used 0.25 Michelson contrast and was repeated at 0.125 contrast. Explicit gray-blank responses were measured for baseline subtraction. Behavior was held at zero.

For each condition, the primary response was the phase-averaged expected count per 1/240-s bin minus the gray-blank response. The preferred motion direction was selected from the acquired responses. At that direction, the complete dynamic SF × TF response surface was fit with the separable and inseparable R0/R1 functions described by Yu et al. [52]; the model was selected by their BIC comparison. The continuous preferred SF and TF were the maximum of the selected fitted surface. Displayed passband contours mark the boundary at 55% of the fitted peak response within the probed frequency range.

The population plots included all 725 matched model units, irrespective of tuning-fit quality. Every unit produced finite preferred-frequency coordinates and a finite passband surface; 723 fits reported optimizer convergence and 2 finite nonconverged fits were included. Recorded gratings did not independently vary temporal frequency, so temporal-frequency values in Figure 4 refer to tuning inferred from the model assay, not directly measured biological TF tuning.

Figure 4E summarizes the population by evaluating every fitted surface on a shared logarithmic SF × TF grid and counting, at each coordinate, the fraction of units whose normalized response exceeded the 55% passband threshold. The displayed grayscale field represents this passband-occupancy fraction.

The two displayed tuning examples were chosen from a stricter subset of 145 units. These units had complete exact-identity assays; phase-converged responses (24 versus 12 phases: surface correlation ≥ 0.995, SF/TF peak shifts ≤ 0.51 octaves, direction shift ≤ 20.1°); positive blank-subtracted drive (≥ 0.002 expected counts/bin); an interior, coherent raw tuning peak; and a successful full-grid Yu fit with *R*^2^ ≥ 0.80 and raw–fit peak discrepancies ≤ 0.75 octaves. At the second contrast, the surface correlation had to remain ≥ 0.80, peak shifts ≤ 0.75 octaves, direction shift ≤ 40.1°, and drive ≥ 0.001 counts/bin. Recorded-neuron SF tuning additionally required at least 50 spikes, split-half correlation ≥ 0.20, a non-boundary peak, and matched-twin SF-curve correlation ≥ 0.50 with peak discrepancy ≤ 1 octave. These gates restrict the illustrative tuning examples, not the 725-unit population analysis.

#### Retinal power redistributed by fixational motion

The distribution-level power analysis used 100 natural-image patches and event-audited one-second epochs from an independently selected candidate bank of 200 fixation records. Windows without sufficient continuous filtering padding were discarded. Natural-image spatial power, *I*(*k*), was estimated from Tukey-windowed Fourier transforms of the retinal image patches and averaged within animal before animals were weighted equally. For each fixation *X*(*t*), retinal translation was represented by its exact unit-modulus carrier

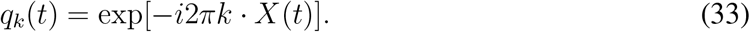

The complete temporal spectrum *Q*(*k, ω*) of this carrier was estimated after temporal demeaning with two discrete prolate spheroidal tapers. The finite observation window and tapers broaden spectral estimates, limiting temporal-frequency resolution. Positive and negative temporal frequencies were folded together and motion directions were averaged. The expected translated-image spectrum was then obtained using the Kuang/Rucci factorization

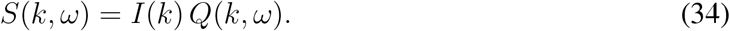

This calculation retains the complete temporal order of each fixation.

For every fixation, the nonzero-TF power map was divided by its integrated dynamic power. Drift windows contained no detected event within the scored interval or 50 ms of either edge. Microsaccade windows contained at least one verified event with 0 *<* amplitude *<* 1°, fully inside the scored interval with 50-ms margins. Both conditions passed the native-validity, event-coverage, and unmatched-motion checks described above. The comparison comprised 47 drift and 18 microsaccade windows. Group assignment was based on eye-movement records, independently of retinal spectra and neural or model responses. For microsaccade-containing windows, the spectra characterize the frequency content of transient retinal motion. Complete trajectories were retained within each window; this comparison is not an additive decomposition into drift and microsaccade motion. Traces were averaged within animal and the two animals were then weighted equally in each condition. Each displayed conditional spectrum therefore integrates to one and shows where dynamic power is distributed rather than how much total power is present.

Translation conserves image power. For each spatial Fourier mode, the TF=0 fraction was the squared magnitude of the mean translation carrier and the TF*>* 0 fraction was its complement; their sum was verified numerically to equal one. Figure 4C shows the drift and microsaccade conditional distributions separately. Figure 4D shows their log_2_ microsaccade-to-drift ratio with the two example passbands overlaid.

#### Direct passband engagement

For Figure 4F, retinal power was computed directly from the 8,000 rendered movies used for the response matrix. Here the estimation window was 250 ms, matching the movies used to predict responses. Each movie underwent a spatial Fourier transform, removal of the temporal mean, and a two-taper temporal spectral estimate with DPSS parameters *NW* = 1.5 and *K* = 2. The resulting power was placed on the measured SF×TF×orientation grid.

For each unit, we computed a weighted average of movie power across spatial frequency, temporal frequency, and orientation, with weights determined by the unit’s grating responses. We constructed these weights by multiplying the fitted SF×TF tuning surface at the unit’s preferred motion direction by its measured orientation weights. To obtain the orientation weights, we used responses at the SF×TF combination with the largest measured response, subtracted the blank response, and set negative values to zero. We then averaged responses across opposite motion directions and normalized the resulting orientation weights to their maximum. The same orientation weights were applied at every SF×TF combination. We normalized the combined nonnegative weights to sum to one, multiplied them by the corresponding movie-power values, and summed across frequencies and orientations. All tuning weights were estimated from the controlled grating assay, independently of the natural-image response outcomes.

For unit *u* and fixation history *h*, we defined tuning engagement as

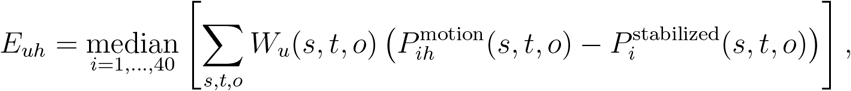

where *i* indexes images, *s, t, o* index spatial frequency, nonzero temporal frequency, and orientation, and *P* denotes the estimated retinal movie power. The nonnegative tuning weights *W*_*u*_ were constructed as described above and normalized so that ∑_*s,t,o*_ *W*_*u*_(*s, t, o*) = 1. For Figure 4F, engagement values were converted to percentiles separately within each unit. We averaged firing rates across matched images separately for the measured-motion and stabilized conditions. SSI was pooled across images using each condition’s expected spike counts as weights. For both measures, we then calculated percent modulation as 100 (*M S*)*/S*, where *M* and *S* denote the pooled measured-motion and stabilized values, respectively. Fixation histories were divided into five within-unit tuning-weighted-power percentile bins. Boxes show the distribution of unit-specific bin medians. The population curve is the median across units, with 95% intervals from 5,000 bootstrap resamples of units and fixation histories within unit.

#### Spatiotemporal tuning engagement compared with movement descriptors

We tested whether engagement improved prediction of rate and SSI modulation beyond movement class and path length using the same 40-image × 200-history response matrix as Figure 4F. The histories comprised 146 drift and 54 microsaccade windows from 187 source trials, with 100 histories from each animal. For each unit and history, engagement was the median motion-induced power weighted by the unit’s spatiotemporal tuning across images. Firing rates were averaged across images, and SSI was pooled across images using expected spike counts as weights, before calculating percent modulation relative to the stabilized condition. We fit separate regressions for rate and SSI modulation in each of the 725 model units, using movement class, path length, engagement, or combinations of these predictors. All regressions included the animal from which the eye-movement history was recorded, and prediction errors were weighted equally between animals. The baseline model included movement class and allowed a separate nonlinear relationship between path length and response modulation for drift and microsaccade histories.

Continuous predictors were transformed to training-set percentiles and represented by five cubic B-spline terms, with interior knots at one-third and two-thirds of the percentile range. All transformations and feature standardization used training data only. Ridge regression used a fixed penalty of 0.01 relative to weighted mean squared loss. We evaluated predictions using five-fold cross-validation, keeping histories from the same source trial together and stratifying folds by animal and movement class. Cross-validation was repeated with three fixed random seeds, and squared prediction errors were averaged across repetitions. For each unit, we calculated the fractional reduction in held-out mean squared error after adding engagement to the movement-class and path-length baseline, then summarized these reductions by their median across units.

#### Cumulative network-stage analysis

To trace the high-passband effect through the trained model, Figure 4G used fixation histories selected solely from the top 20% of each unit’s passband-power distribution. 10 histories spanning the high-engagement range were crossed with 10 natural images, producing 100 measured/stabilized movie pairs. Unit effects were summarized from the selected histories lying within that unit’s top quintile.

The trained readout logits were partitioned according to the feature channels contributed by the first, second, and third spatial stages. The fixed zero-behavior contribution and readout bias were included once. Cumulative rate maps are formed from (1) the first spatial stage, (2) the first and second spatial stages, and (3) all three spatial stages, summing every readout factor associated with each included channel group. The model’s softplus was applied to each cumulative logit. We verified numerically that the final cumulative map reproduced the full model output.

To remove trivial uniform gain, temporal response modulation was measured as

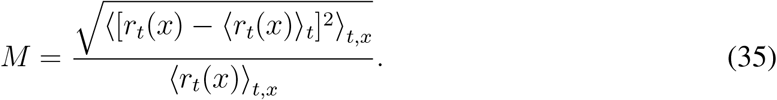

Figure 4G reports 100(*M*_motion_ − *M*_stabilized_) in percentage points. Spatial sharpening was the expected-spike-weighted absolute difference *I*_motion −_ *I*_stabilized_ in bits per spike. Unit effects were first summarized across the high-passband histories and then pooled by their population median; intervals were obtained from 5,000 bootstrap resamples of units.

For the zero-improvement tests in Figure 4B,F,G, we retained each panel’s population estimand and paired bootstrap scheme described above. Two-sided bootstrap sign-tail probabilities were computed as *p* = min {1, 2[1 + min(*n*_≤0_, *n*_≥0_)]/(*N*_boot_ + 1)}, where *n*_≤0_ and *n*_≥0_ count resampled population effects on either side of zero. We applied Holm correction jointly across all 32 bin- or stage-specific tests (two outcomes in eight path-length bins, five engagement quintiles, and three cumulative readouts). These tests assess the population summaries under the stated resampling schemes, conditional on the fitted model; the boxplots separately describe heterogeneity across units.

## Data and code availability

The data and associated analysis and modeling code will be made available upon publication.

## Acknowledgments

This work was supported by NEI K99-EY037839-01 to D.P.R., NEI R01-EY030998 to J.F.M., NEI R01-EY018363 to M.R., Fuster Endowment funds to A.C.H., and NEI K99/R00-EY032179 to J.L.Y. We thank Yonatan Abrham (Mitchell Lab) for collecting some of the data used in this report.

## Supplementary Information

### Supplemental figures

**Extended Data Figure 1.**
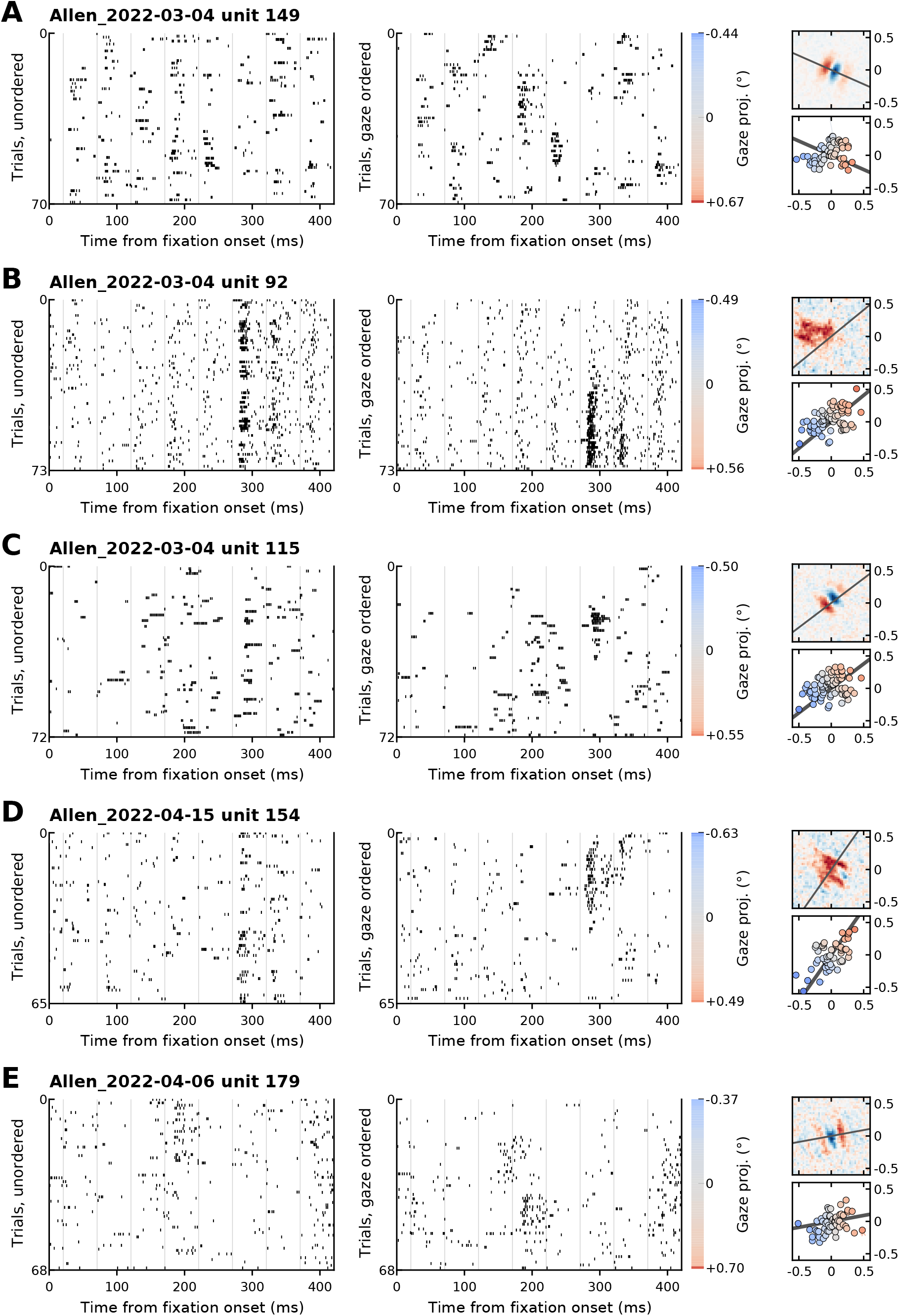
Eye-position sorting reveals single-cell response structure across example V1 units. Additional single-cell examples of the gaze-dependent trial structure shown in Fig. 1D–G. Panel **(A)** shows the same unit as Fig. 1D–G, unit 149 from Allen 2022-03-04, and includes the unordered trial raster for comparison. In each panel, left, trial rasters aligned to fixation onset with trials shown in their original unordered arrangement. Middle, the same responses reordered by gaze position projected onto the axis of maximal spatial sensitivity for that unit. Color strip, signed gaze projection for the gaze-ordered trials, with blue and red indicating opposite positions along the projection axis. Right, spike-triggered average and gaze positions for the sorting window; the black line indicates the projection axis used for ordering trials. Across units, ordering trials by eye position reveals response structure that is less apparent in the unordered rasters, consistent with fixational eye movements contributing to single-trial variability in foveal V1.

**Extended Data Figure 2.**
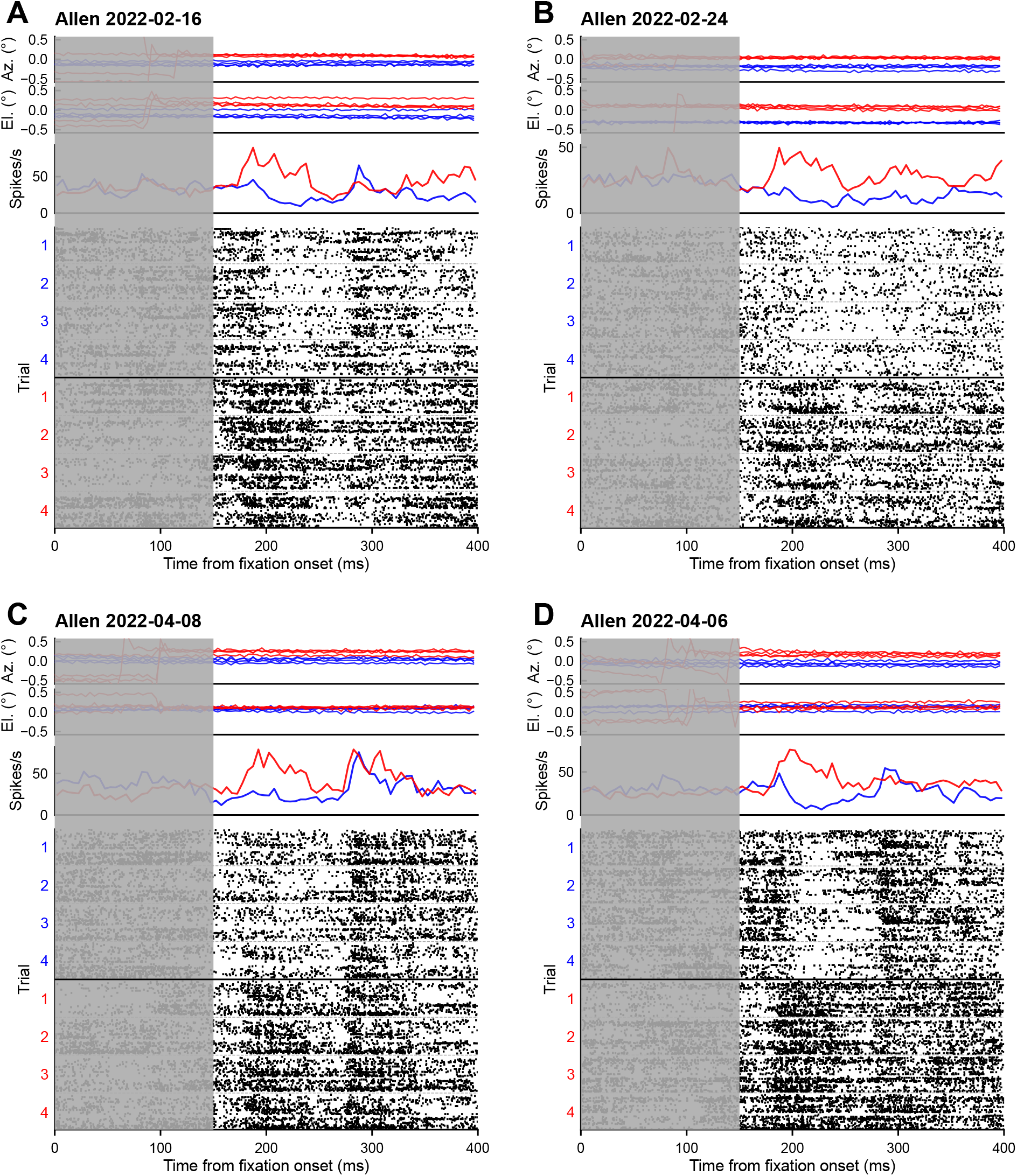
Gaze-dependent population structure across recording sessions. Additional examples of the population-level effect shown in Fig. 1H–J. Panels **(A)–(D)** show four sessions from monkey Allen. Top, horizontal and vertical gaze traces for two groups of four trials, colored blue and red; gray shading marks an initial context interval. Middle, population-averaged spike rate for the same trial groups. Bottom, population rasters for the same trials, with units stacked within trials and the black line separating blue and red groups. Across sessions, trials with similar gaze trajectories show similar population activity, whereas groups with distinct gaze trajectories show systematic response differences.

**Extended Data Figure 3.**
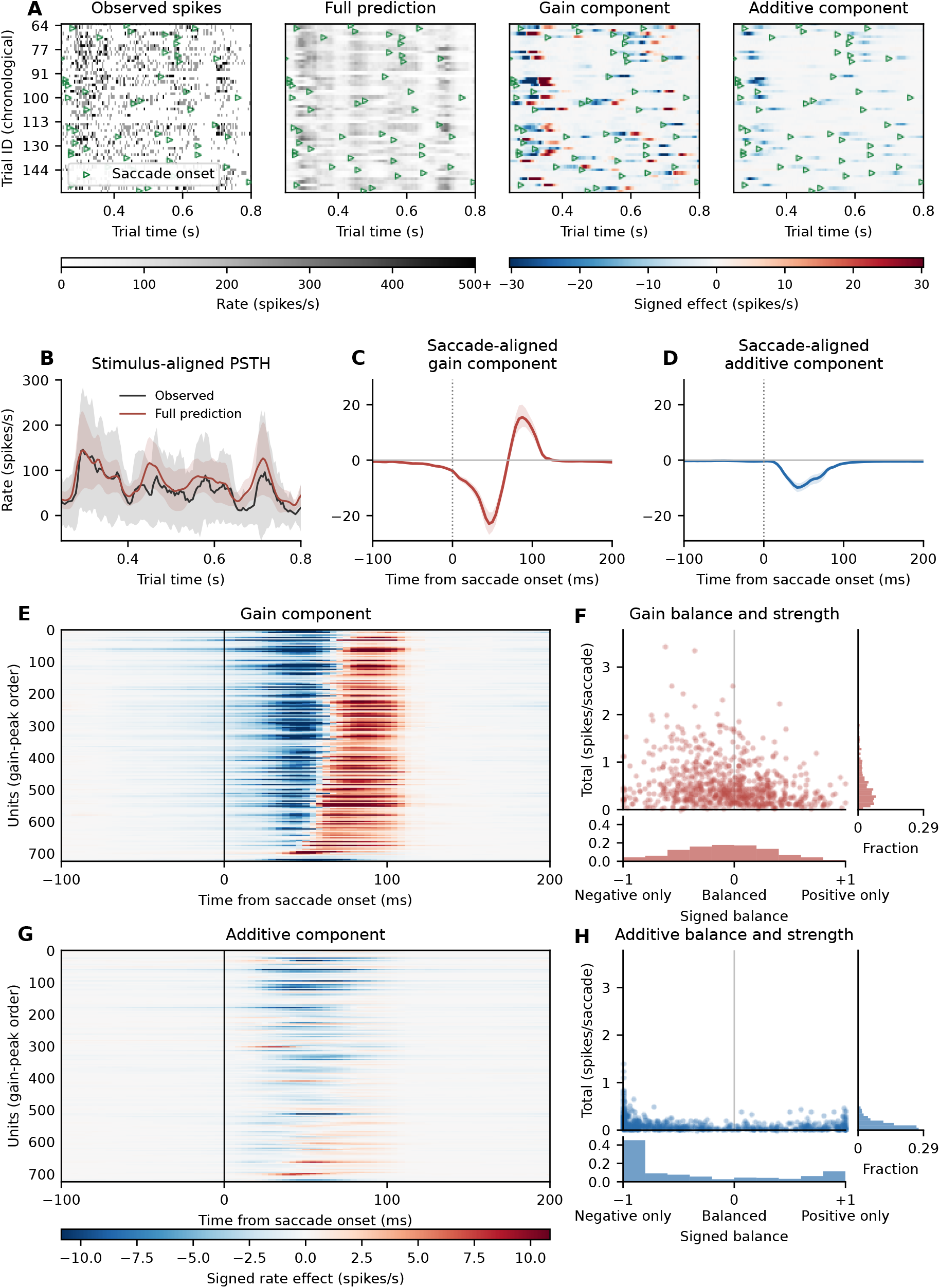
The model’s extraretinal pathway learns structured saccade-linked modulation. Despite its small contribution to overall prediction performance during the flashed image experiment (Fig. 3), the extraretinal pathway produces consistent peri-saccadic changes in predicted firing rate. The pathway learns arbitrary transformations of eye position and velocity without being given explicit saccade labels. Its learned effects show a consistent saccade-related pattern, with suppression followed by enhancement in the gain component and predominantly monophasic suppression or enhancement in the additive component. **(A)** From left to right, observed responses, full-model predictions, and rate effects of the gain and additive components for an example unit across 56 repeated stimulus presentations. Green triangles mark detected saccade onsets. To isolate each component’s effect, we first calculated the model’s internal drive in the absence of extraretinal modulation. We then restored the eye-dependent change in either the gain or additive component, holding the other unchanged. Each rate effect is the difference between this reconstructed prediction and the prediction without extraretinal modulation, evaluated after the output nonlinearity and converted to spikes/s. All predictions retain the same recorded retinal input and fitted weights. Because of the output nonlinearity, the two component effects need not sum to the full extraretinal rate change. **(B)** Observed and full-model stimulus-aligned mean responses for the trials shown in A, smoothed over three bins (12.5 ms). Shading indicates ±1 standard deviation across trials. **(C)** Saccade-aligned rate effect of the gain component for the example unit, showing suppression followed by enhancement. Shading indicates 95% bootstrap intervals from 400 resamples of trials. **(D)** Saccade-aligned rate effect of the additive component for the same unit, showing predominantly suppressive modulation. Shading as in C. **(E)** Saccade-aligned gain-component rate effects across the 725 units analysed in the main manuscript text. Each row shows one unit’s mean rate effect across saccades. Rows are ordered by the time of peak enhancement within 200 ms after saccade onset, with later peaks above earlier peaks and units without a positive peak at the bottom. Suppression followed by enhancement is common across the population. **(F)** Balance and magnitude of gain modulation for each unit. Enhancing area *P* and suppressive area *N* were calculated by integrating the positive and absolute negative portions of each unit’s saccade-averaged rate effect from 0 to 200 ms. Total modulation (vertical axis), *P* + *N*, is expressed in spikes/saccade. Signed balance (horizontal axis), (*P − N*)/(*P* + *N*), ranges from −1 for purely suppressive effects to +1 for purely enhancing effects, with zero indicating equal areas, as in a balanced biphasic response. Marginal histograms show the fraction of units in each bin. Gain effects concentrate near balanced enhancing and suppressive contributions (median absolute signed balance, 0.296), with a median total modulation of 0.406 spikes/saccade. **(G)** Saccade-aligned additive-component rate effects, using the same units, ordering, and color scale as E. Effects are generally monophasic and predominantly suppressive, with enhancement in a subset of units. **(H)** Balance and magnitude of additive modulation, calculated and displayed as in F. Signed balances concentrate toward the extremes, particularly the suppressive end (median absolute signed balance, 0.869), with a median total modulation of 0.092 spikes/saccade. Together, these analyses show that the model’s extraretinal pathway produces structured peri-saccadic modulation despite its small contribution to overall prediction performance during the flashed-image experiment. The model could learn arbitrary transformations of eye position and velocity, without explicit saccade labels or a prescribed temporal response. Nevertheless, it learned a consistent saccade-linked pattern across units, with suppression followed by enhancement in the gain component and generally monophasic suppression or enhancement in the additive component. The biphasic time course qualitatively resembles the extraretinal modulation reported in macaque V1 by McFarland et al. [53], although the decomposition into gain and additive effects differs between the studies. These findings are consistent with a saccade-linked extraretinal contribution in our data.

**Extended Data Figure 4.**
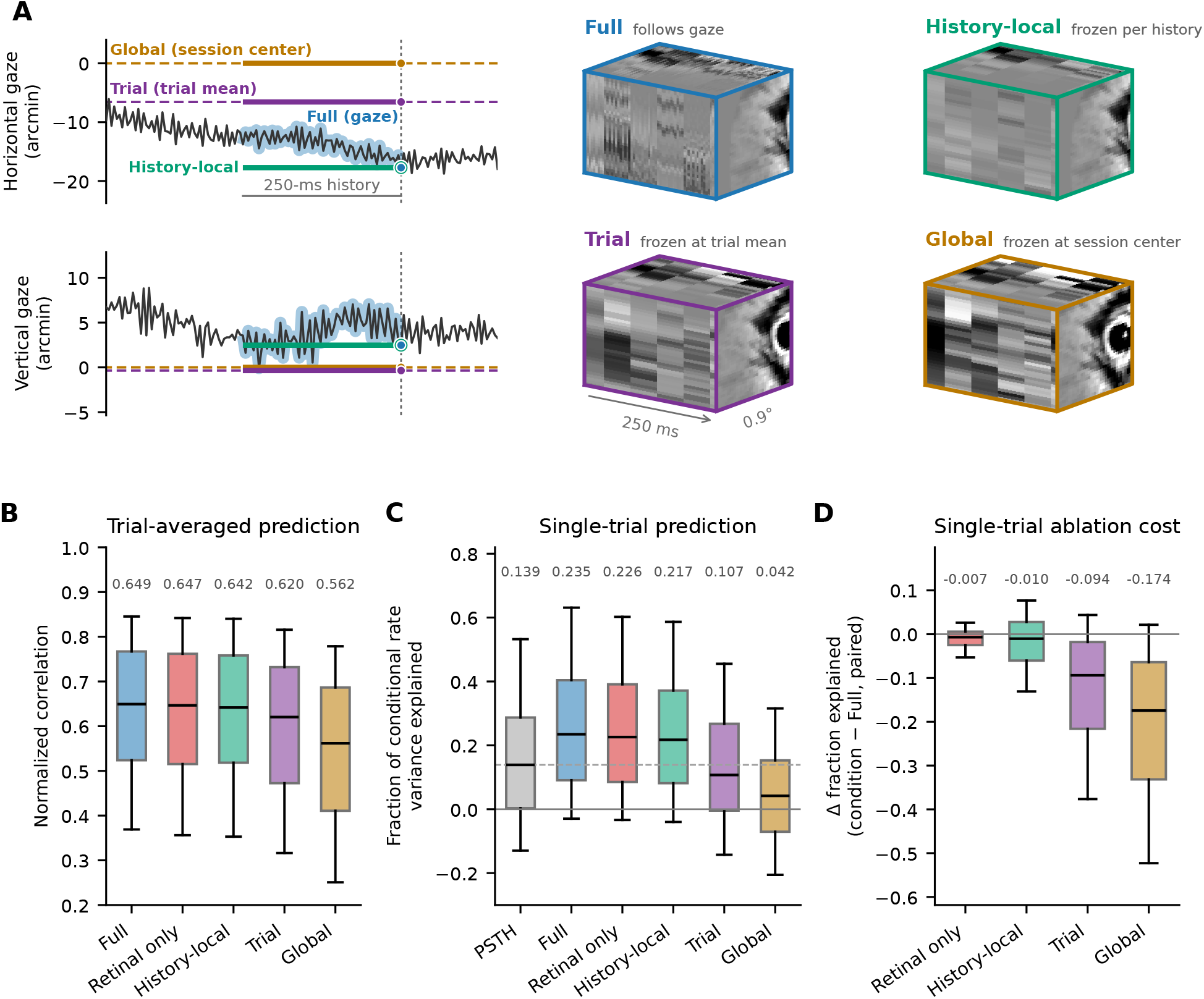
Prediction accuracy declines as retinal stabilization extends from input histories to whole trials and sessions. All conditions use the Figure 3 model with fixed weights and the same scored trials and units as Figure 3C,D (*n* = 962 units across 19 sessions). **(A)** Retinal input under each stabilization condition, illustrated for one 250-ms input history. The left plots show horizontal and vertical gaze position relative to the session center. The black trace shows measured gaze, the dashed vertical line marks the time of the prediction, and the bar marks the 250-ms history that supplies the model input. Colored lines show the gaze positions that are used to render the 60 input frames for each condition. Full (blue) follows measured gaze. History-local (green) holds gaze at its latest position in the history, so the final retinal frame matches Full while preceding gaze displacement is removed. Trial (purple) holds gaze near the trial’s mean position. Global (ochre), the stabilized-retina condition of Figure 3, holds gaze at one session-wide reference. The right plots show the retinal input that results under each condition (0.9°× 0.9° × 250 ms). All four conditions retain the extraretinal inputs and the 20-Hz flashed-image sequence. Retinal only (B–D) is the retinal-only condition of Figure 3. **(B)** Trial-averaged prediction, measured as noise-normalized correlation. Median paired changes were −0.002 for History-local minus Full (95% CI [−0.009, +0.002]), −0.024 for Trial minus History-local ([− 0.038, 0.013]), and −0.045 for Global minus Trial ([− 0.056, −0.035]). **(C)** Single-trial prediction, measured as the fraction of conditional rate variance explained with the Figure 3D denominator. The dashed line marks the PSTH median and the solid line marks zero explained variance. **(D)** Change in single-trial prediction relative to Full, calculated within each unit before pooling. Median changes were− 0.007 for Retinal only (95% CI [−0.011, −0.004]), 0.010 for History-local ([−0.025, −0.004]), −0.094 for Trial ([−0.131, −0.068]), and −0.174 for Global ([−0.203,−0.152]). Between successive stabilization conditions, median paired changes were −0.079 for Trial minus History-local ([−0.111, −0.057]) and −0.066 for Global minus Trial ([−0.085, −0.044]). The within-unit difference between the History-local and Retinal-only changes was 0.001 ([−0.007, +0.012]), which did not establish a larger cost for removing recent gaze displacement than for removing the extraretinal inputs. Preserving each history’s final retinal position avoided most of the loss caused by stabilization across whole trials or sessions, with a smaller contribution from gaze displacement within the history. We note that the 20-Hz flashed-image sequence provided sustained temporal drive even when gaze was held fixed. Recent retinal motion is therefore expected to contribute more under naturalistic conditions without rapid image flashes, as explored in the model analyses of Figure 4. Boxes span the interquartile range, with median lines and 10th–90th percentile whiskers. Medians are displayed above the box plots in B–D. The 95% confidence intervals were calculated using 10,000 paired session-bootstrap samples.

